# AMPKα1 is essential for Glucocorticoid Receptor triggered anti-inflammatory macrophage activation

**DOI:** 10.1101/2020.01.02.892836

**Authors:** Giorgio Caratti, Thibaut Desgeorges, Gaëtan Juban, Mascha Koenen, Bozhena Kozak, Marine Théret, Bénédicte Chazaud, Jan P Tuckermann, Rémi Mounier

**Affiliations:** Institute of Comparative Molecular Endocrinology, 8/1 Helmholzstraße, Universität Ulm, Ulm, Germany; Institut NeuroMyoGène, Université Claude Bernard Lyon 1, CNRS UMR 5310, INSERM U1217, Université Lyon, Lyon 69008, France

## Abstract

Macrophages are key immune cells which mediate both the acute inflammatory phase and the repair phase after tissue damage. Macrophages switch from pro-inflammatory to anti-inflammatory cells that sustain repair and return to tissue homeostasis. We show that the metabolic sensor, AMP-activated protein kinase (AMPK) is essential for glucocorticoid induction of an anti-inflammatory macrophage phenotype. While canonical gene regulation by glucocorticoids was not affected by loss of AMPK, we identified AMPK-dependent glucocorticoid-regulated genes in macrophages, related to efferocytosis. AMPK-deficient macrophages do not acquire phenotypic and functional anti-inflammatory features upon glucocorticoid exposure. We identified FOXO3 as an AMPK-dependent regulator of glucocorticoid activity in macrophages. Loss of AMPK in macrophages *in vivo* abrogates glucocorticoid anti-inflammatory actions during post-injury muscle regeneration and endotoxin induced acute lung injury. These data highlight that the glucocorticoid receptor is dependent on AMPK for its immunomodulatory actions in macrophages, linking their metabolic status to transcriptional control in resolving inflammation.

## Introduction

Resolution of inflammation is essential for the return to tissue homeostasis after an inflammatory insult, both to limit excessive organ damage, but also to initiate the repair process (Watanabe et al., 2019). Resolution involves the suppression of pro-inflammatory active cells and the acquisition of an anti-inflammatory/restorative phenotype by immune cells. These together drive the switch from bactericidal activity and cytokine release to tissue repair. Macrophages are central to this process, being involved in both the acute inflammatory phase and the repair phase (Watanabe et al., 2019; Wynn and Vannella, 2016). Inflammatory macrophages rapidly infiltrate the target tissue, drawn by chemokines such as CCL2 or CX3CL1, and release pro-inflammatory cytokines such as TNF-α and CCL3 (Mantovani et al., 2004; Turner et al., 2014). During the inflammatory phase, macrophages then phagocytose apoptotic cells in a process called efferocytosis. Engulfment of apoptotic cells provokes a macrophage phenotypic switch to an alternative, anti-inflammatory, pro-repair state, which promotes tissue homeostasis (Ortega-Gomez et al., 2013). Similarly, anti-inflammatory signals can push macrophages towards a pro-efferocytosis phenotype (Fadok et al., 1998; Nepal et al., 2019; Weigert et al., 2006) assisting in the regenerative function of macrophages. Glucocorticoids (GCs) are one of the most potent immuno-suppressive agents, which promote efferocytosis in macrophages (Desgeorges et al., 2019a; McColl et al., 2009). GCs are also the most frequently prescribed anti-inflammatory agents, with over 1% of the UK, Danish and American populations, at one point, having received a prescription for GCs (Fardet et al., 2011; Laugesen et al., 2017; Overman et al., 2013). GCs signal through the glucocorticoid receptor (GR), a member of the nuclear receptor superfamily, which acts as a transcription factor to regulate gene expression upon exposure to ligand (Weikum et al., 2017b). Unlike many other nuclear receptors, GR resides in the cytoplasm, where it can interact with various signaling enzymes (Petta et al., 2016). Upon ligand exposure, GR translocates to the nucleus to mediate its effects (Vandevyver et al., 2012a) regulating transcription through transactivation and transrepression (Weikum et al., 2017b). Suppression of pro-inflammatory cytokines occurs at the transcriptional level through GR tethering to pro-inflammatory transcription factors such as NFκB and AP-1, or by direct binding to DNA at, or nearby NFκB and AP-1 response elements and represses transcription through a tethering independent mechanism (Hudson et al., 2018; Uhlenhaut et al., 2013; Weikum et al., 2017a). Suppression of cytokines was considered as the canonical mechanism of immunosuppression by GCs for decades (Barnes, 2011). However, recent studies implicated that GR dependent upregulation of anti-inflammatory genes plays an essential role in the resolution of inflammation. In particular the expression of *Tsc22d3 (Gilz), Dusp1, Sphk1, Ikkb* and *A20* (Eddleston et al., 2007; Oh et al., 2017; Vandevyver et al., 2012b; Vettorazzi et al., 2015) are important for the anti-inflammatory properties of GCs. Little is known however, regarding how GR cross-talks with other key regulators of the anti-inflammatory status of macrophages. One such regulator is AMP-activated protein kinase (AMPK), which controls a wide variety of metabolic functions in all cell types (Hardie et al., 2012). AMPK is activated by a high AMP:ATP ratio or through diminished cellular glucose concentration (Hardie et al., 2012), making AMPK a sensor of cellular energy deprivation. Upon activation, AMPK phosphorylates various proteins to inhibit cellular anabolism and activate catabolism (Hardie et al., 2012). For example, the fatty acid elongation enzyme, acetyl-CoA carboxylase, is phosphorylated and inhibited in response to AMPK activation, preventing the anabolic process of fatty acid elongation. AMPK plays a decisive role in the promotion of anti-inflammatory macrophages, initiating the repair phenotype (Mounier et al., 2013; Sag et al., 2008). However, the mechanisms by which AMPK signaling contributes to other anti-inflammatory signals is not understood. GR and AMPK were previously shown to communicate in their role as regulators of metabolism in non-immune cells (Christ-Crain et al., 2008; Nader et al., 2010; Ratman et al., 2016; Seelig et al., 2017). We therefore sought to determine whether AMPK regulates GR anti-inflammatory actions and controls the tissue regenerative actions of macrophages.

We describe a functional interaction between GR and AMPK signaling, whereby deletion of AMPKα1 in macrophages alters GR mediated gene regulation resulting in loss of regulation of a subset of GR target genes, dubbed non-canonical, but maintenance of the regulation of traditional GR target genes. These non-canonical genes are enriched for FOXO3 targets, and we identified an AMPK-dependent loading of FOXO3 together with GR at their regulatory regions. We demonstrate that GR requires AMPK activity for the development of anti-inflammatory macrophages while leaving canonical suppression of cytokine signaling intact. Loss of AMPK abrogates GC induced efferocytosis *in vitro*, limiting the potential anti-inflammatory actions of GCs. Furthermore, we demonstrate that detachment of the signaling between GR and AMPK *in vivo*, using a myeloid specific deletion of AMPKα1, impairs GC-driven resolution of tissue repair (i.e. skeletal muscle injury) and acute lung inflammation induced by endotoxin shock.

## Results

### The glucocorticoid receptor signals through AMPKα1 in macrophages

GCs and AMPK activators both exert anti-inflammatory actions on macrophages (Arnold et al., 2007; Ehrchen et al., 2007; Mounier et al., 2013; Xue et al., 2014). Therefore, we sought to determine whether AMPK and GR signaling converge in macrophages. We first investigated whether the GR agonist dexamethasone (Dex) activates AMPK in bone marrow derived macrophages (BMDMs). BMDMs were treated with the AMPK activating compound 991, Dex or a combination of both and AMPK signaling was assessed. After one hour of treatment, Dex increased the phosphorylation of AMPK at T172 as did 991, but with no additive effect of co-treatment of Dex and 991 (Fig.1A,B). No effect was observed at 30 min of treatment by any compound (Fig.S1A). The induced phosphorylation also resulted in increased activity of AMPK, as the downstream target ACC was also phosphorylated in response to 1 hour Dex treatment (+32%) (Fig.1A,C). Thus, Dex activates AMPK in macrophages in a similar extent than pharmacological activators after 1 hour.

**Figure 1.**
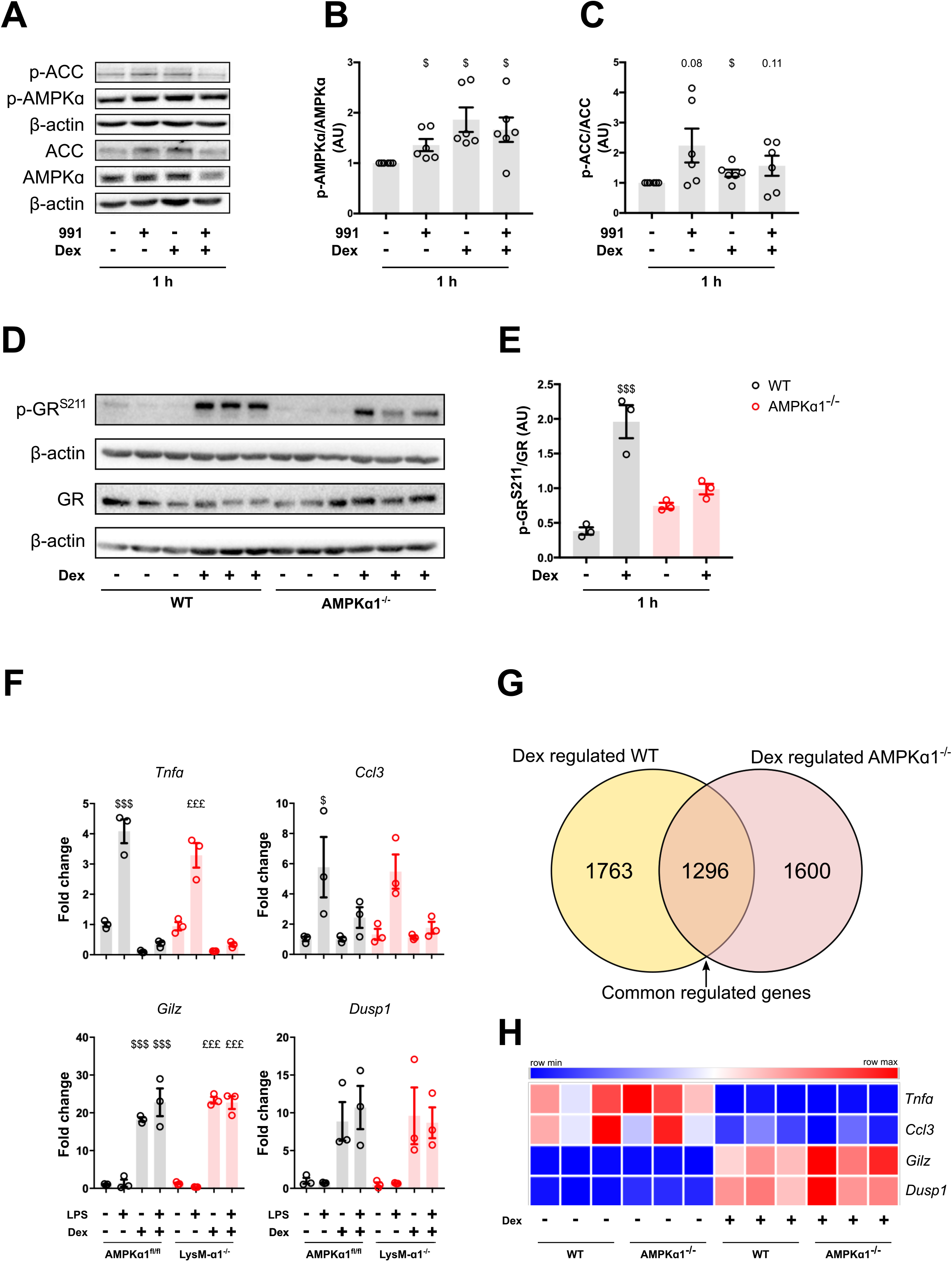
Glucocorticoid receptor signaling *via* AMPKa1 in macrophages. **(A-C)** Bone marrow-derived macrophages (BMDMs) were treated with 991 and with the glucocorticoid (GC) dexamethasone (Dex) for 60 minutes and the phosphorylation of the AMPK target Acetyl CoA Carboxylase (ACC) on Ser79 and the phosphorylation of Thr172 of AMPKα were assessed by immunoblot. Quantification of P-AMPK **(B)** and of P-ACC **(C)** shows ratio between the phospho-protein on total-protein both normalized on β-actin. **(D-E)** BMDMs from WT and AMPKα1^−/−^ mice were treated with Dex for 60 minutes and glucocorticoid receptor (GR) phosphorylation at Ser211 was evaluated by immunoblot. Quantification of GR^S211^ phosphorylation **(E)** shows ratio between the phospho-protein on total-protein normalized to loading control. **(F-H)** BMDMs from WT and AMPKα1^−/−^ mice, or AMPKa1^fl/fl^ and AMPKa1^fl/fl^;LysM^Cre/+^ (LysM-α1^−/−^) were analyzed by RT-qPCR or RNA-seq. **(F)** RT-qPCR was performed on a series of GR target genes in AMPKa1^fl/fl^ or LysM-α1^−/−^ BMDMs treated with vehicle, Dex, LPS or LPS+Dex for 24 hours. **(G)** Venn diagram of the numbers of genes regulated by Dex in WT and AMPKa1^−/−^ macrophages from RNA-seq. **(H)** Heat map from RNA-seq data showing the expression of specific GR target genes. Results are means ± SEMs of 6 (A-C) or 3 (D-H) experiments. $ p<0.05, $$$ p<0.001 *vs* untreated WT; £ p<0.05, £££ p<0.001 *vs* untreated LysM-α1^−/−^/AMPKα1^−/−^, using Student’s t test (A-C) or one-way ANOVA (D-E-F).

Previous reports indicate that AMPK activation may induce GR phosphorylation and activation through p38, at least in the liver (Nader et al., 2010). Therefore, we determined how genetic loss of AMPKα1, the only AMPK catalytic subunit expressed in macrophages (Sag et al., 2008), affects GR phosphorylation. Both wildtype (WT) and AMPKα1 knockout (AMPKα1^−/−^) BMDMs were treated with Dex for 1 hour, and phosphorylation of GR at S211 was assessed by immunoblot. Upon Dex treatment, GR phosphorylation was increased (+511%), however in macrophages lacking AMPK this increase was strongly diminished (−49%) (Fig.1D,E). In these conditions, we did not observe any effect on p38 activation (determined through p38 phosphorylation at T180/Y182) (Fig.S1B). This suggests that p38 activity is independent of AMPK status and does not correlate with GR^S211^ phosphorylation in macrophages. Since GR^S211^ phosphorylation is associated with activity of GR in the nucleus (Chen et al., 2008), we tested whether AMPK loss affected GR translocation to the nucleus. We did not see any effect on GR translocation indicating that loss of AMPK does not affect GR trafficking in macrophages (Fig.S1C,D).

The main mode of action of GR is as a transcriptional regulator. Therefore, we sought to determine how loss of AMPK alters the gene regulatory activity of GR. To our surprise, despite showing previously attenuated S211 phosphorylation of GR upon loss of AMPKα1, we did not observe altered regulation of a panel of classically regulated index genes encoding anti-inflammatory proteins and inflammatory mediators by GR. The transactivated genes (*Gilz*, *Dusp1*), and the transrepressed genes (*Tnf*, *Ccl3*) in BMDMs treated with vehicle, Dex, LPS (lipopolysaccharide) or LPS+Dex for 24 hours were all regulated in the same manner in both WT (LysM^+/+^;AMPKα1^fl/fl^) and LysM^Cre/+^;AMPKα1^fl/fl^, (hereafter indicated as LysM-α1^−/−^) macrophages when assessed by RT-qPCR (Fig.1F). A panel of genes assessed after 6 hours of treatment were similarly regulated (Fig.S1E) indicating no time-dependent effect of AMPK on GR activity. As there was no obvious effect on classically GC-regulated genes, we performed RNA expression profiling on WT and AMPKα1^−/−^ macrophages using RNA-seq to identify potentially novel GR-AMPK regulated genes (Fig.1G, Supplementary Table 1). Differentially expressed genes between WT and AMPKα1^−/−^ were identified using a p<0.05 as a cut off. We identified 3059 and 2896 genes regulated by Dex in WT and in AMPKα1^−/−^ BMDMs, respectively, 1296 genes being found commonly regulated (Fig.1G, Supplementary Table 1). Within this overlap of 1296 genes, we confirmed that *Tnf*, *Ccl3*, *Gilz* and *Dusp1* were similarly regulated by Dex in both WT and AMPKα1^−/−^ BMDMs (Fig.1H), confirming the RT-qPCR results (Fig.1F, S1E). However, a substantial fraction of Dex-regulated genes was differentially expressed exclusively in WT BMDMs (1763) and in AMPKα1^−/−^ deficient (1600) macrophages, supporting the need of AMPK to modulate Dex-triggered gene expression. Gene ontological analysis of the AMPK-dependent transcriptome (i.e. WT selective) in response to GC treatment was highly enriched in genes involved in protein deubiquitination but also mitochondrial function, while the AMPKα1^−/−^ selective response to GCs revolved around up regulation of protein synthesis, likely due to loss of AMPK-dependent suppression of mTOR activity, but also negative regulation of cell proliferation (Fig.S1F).

### AMPKα1 in macrophages is required for glucocorticoid protection during acute lung inflammation and endotoxin shock

Having established that loss of AMPKα1 alters GC-dependent transcriptome in macrophages, we aimed to establish functional consequence for the dependence of AMPKα1 on GC signaling. Therefore, we used two mouse models of inflammation, the skeletal muscle regeneration (representative of sterile inflammation) and the lethal endotoxin shock (mimicking systemic inflammation upon bacterial challenge). To assess how GR-AMPK cross-talk in macrophages regulates regeneration of skeletal muscle *in vivo*, we used a tissue specific knockout of AMPKα1 in the myeloid lineage (LysM-α1^−/−^). Macrophages isolated from LysM-α1^−/−^ mice respond to Dex in a similar way as cells from global knockout mice (AMPKα1^−/−^) *in vitro* (Fig.1G and Fig.1I) making them an ideal model for *in vivo* investigation of GR-AMPK cross-talk. We therefore treated WT and LysM-α1^−/−^ mice with vehicle, 10 mg/kg LPS or LPS + 1 mg/kg Dex, before an intravenous injection of oleic acid to induce lung inflammation (Vettorazzi et al., 2015). After 24 hours, mice were sacrificed and cells infiltrating into the lungs were counted in the broncho-alveolar lavage (BAL) fluid. LPS increased the number of cells in the BAL of both WT and LysM-α1^−/−^ mice (4 and 5-fold, respectively) (Fig.2A). Strikingly, the repressive effects of Dex on the reduction of cell numbers in BAL were only observed in the WT, but not in mice lacking AMPK in myeloid cells. Furthermore, we detected fewer pro-inflammatory macrophages in the WT mice after Dex treatment (−70%), but LysM-α1^−/−^ mice were unaffected (Fig.2B,C). Strikingly according to the effects observed *ex vivo*, the effects of GCs were maintained on cytokine production, as we saw no effect on BAL cytokine content upon AMPKα1 deletion (Fig.2D, Fig.S2A). INFγ, TNFα, IL-6 and IL-10 (among others) were all increased in mice treated with LPS, and reduced by Dex treatment in both WT and LysM-α1^−/−^ mice (Fig.2D, Fig.S2A). Nevertheless, at the tissue level, LPS treatment induced alveolar collapse in both WT and LysM-α1^−/−^ mice, which was partially recovered in WT mice treated with Dex, but not in LysM-α1^−/−^ mice (Fig.2E). Both tissue damage and cytokine production are related to death during sepsis (Stearns-Kurosawa et al., 2011). Therefore, we determined whether loss of AMPK in macrophages altered GC effects on survival. We did not see a difference in survival after 50 hours in WT or LysM-α1^−/−^ mice treated with LPS alone (Fig.2F). However, when mice were treated with both LPS and Dex, we found that LysM-α1^−/−^ mice were more likely to die than WT mice (Fig.2G). However, we did not see a genotype dependent effect on temperature or weight loss, likely due to temperature and weight reductions being the main criteria for sacrifice during sepsis (Fig.S2B,C). These data demonstrate that GCs suppressed inflammation during acute lung injury through AMPK in macrophages to reduce the number of pro-inflammatory macrophages, improve tissue recovery and survival.

**Figure 2.**
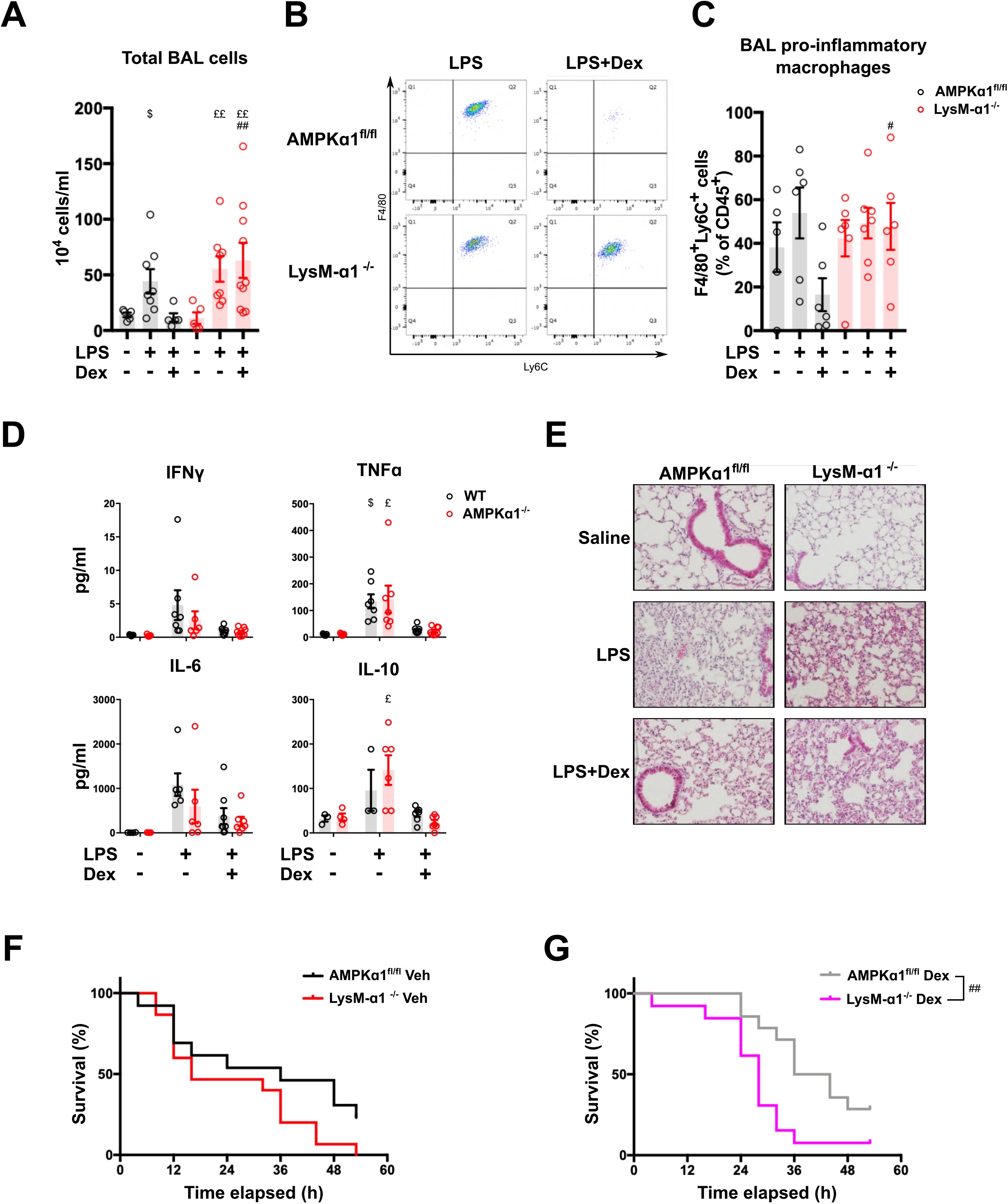
AMPKα1 in macrophages is required for glucocorticoid dependent resolution of inflammation during acute lung inflammation and endotoxin shock. AMPKα1^fl/fl^ (WT) and LysM-α1^−/−^ mice were treated with vehicle or with 10 mg/kg LPS or LPS + dexamethasone (Dex, 1 mg/kg), before an intravenous injection of oleic acid to induce lung inflammation. **(A-C)** After 24 hours, mice were culled and cells infiltrating into the lungs were analyzed. **(A)** The number of cells present in the broncho-alveolar lavage (BAL) fluid was counted. **(B-C)** The number of pro-inflammatory macrophages (F4/80^pos^ Ly6C^pos^) was evaluated by flow cytometry (representative dotplot in **B**) as a percentage of CD45^pos^ cells **(C)**. **(D)** The BAL cytokine content was measured by Luminex after 24 hours. **(E)** Hematoxylin-eosin staining of lung tissue after 24 hours. **(F-G)** Survival curve of mice treated with vehicle **(F)** or Dex **(G)**. Results are means ± SEMs of 5-10 (A), 5-7 (B), 3-7, Data below detection threshold were excluded from analysis (C,D), or 13-15 (F,G) animals. $ p<0.05 *vs* untreated WT; ££ p<0.01 *vs* untreated LysM-α1^−/−^/AMPKα1^−/−^; # p<0.05, ## p<0.01, compared KO *vs* WT for each condition, using Kruskal Wallis (A), ANOVA (C, D) and Gehan-Breslow-Wilcoxon (F,G) tests.

### AMPKα1 in macrophages is required for the glucocorticoid-induced effects on post-injury muscle regeneration

Due to the striking effect we observed on GC action in a model of acute lung inflammation, we next aimed to determine whether the anti-inflammatory actions of GCs could promote muscle repair, despite the usual catabolic effects of GCs. To determine the most effective time point of GC treatment to promote muscle repair, both WT and LysM-α1^−/−^ mice were injected in the *Tibialis Anterior* muscle with cardiotoxin (CTX) to induce muscle damage, a model in which the resolution of inflammation occurs around day 2 (Mounier et al., 2013; Varga et al., 2016a). When Dex (0.05 mg/kg) was given from day 0 or 1 or 2 after injury (*i.e.* before and during the resolution of inflammation) until day 7, muscle regeneration was strongly impaired in WT muscles, as shown by the remaining inflammation and necrotic myofibers at day 8 as compared with PBS treated muscle injured animals (Fig.S3A), confirming that the initial pro-inflammatory phase, preceding the anti-inflammatory repair phase, must not be shortened for an efficient muscle regeneration (Perdiguero et al., 2011). When Dex was administered from day 3 to day 7 after injury at 0.1 mg/kg, WT muscles exhibited a higher number of small myofibers than control muscles 14 days after injury (Fig.S3B). This suggests that daily exposure to Dex, even at low doses, induced myofiber atrophy, which is a well-known GC side effect (Schakman et al., 2009). We then injected a single dose of Dex at day 3 after injury. Here again, injection of a single 1 mg/kg GC dose induced myofiber atrophy (Fig.S3C). We therefore used a single 0.1 mg/kg GC dose at day 3 after injury, which did not cause any detrimental reduction in myofiber size in WT animals (Fig.S3D), and we analyzed GC effects on the maturation of myofibers, which is an indicator of the regeneration process (Fig.3A). Sarcoplasm contains myosin heavy chains (MHC), which compose the contractile apparatus. During adult skeletal muscle regeneration, the first MHC expressed in new regenerating myofibers is the embryonic form of MHC (embMHC). Kinetics of embMHC expression in untreated WT muscle indicated a strong decrease (from 59.3 to 5.6% between day 8 and day 14) of myofibers expressing embMHC, indicative of the maturation of the new regenerating myofibers (Fig.3B,C). Dex treatment decreased the number of embMHC^pos^ myofibers at day 8 in WT mice (−23%), suggesting that Dex accelerated myofiber maturation (Fig.3B,C). As compared with WT, untreated LysM-α1^−/−^ muscle showed a 2-fold increase of embMHC^pos^ myofibers at day 14, indicative of a delay in the maturation of regenerating myofibers (Fig.3B,C) in accordance with the defect of regeneration previously shown in that genotype (Mounier et al., 2013). We did not find any alteration of embMHC expression in Dex treated LysM-α1^−/−^ muscles as compared with vehicle treated LysM-α1^−/−^ muscles (Fig.3B,C), suggesting that Dex had no effect on myofiber maturation when AMPKα1 was absent in macrophages. These data show that a single Dex injection accelerated the maturation of regenerating myofibers in a macrophage AMPKα1-dependent manner. Finally, Two weeks after injury, Dex did not induce muscle atrophy in WT mice, whereas Dex decreased mass of LysM-α1^−/−^ muscles showing that GCs impaired skeletal muscle regeneration in absence of AMPKα1 in macrophages (Fig3.D).

**Figure 3.**
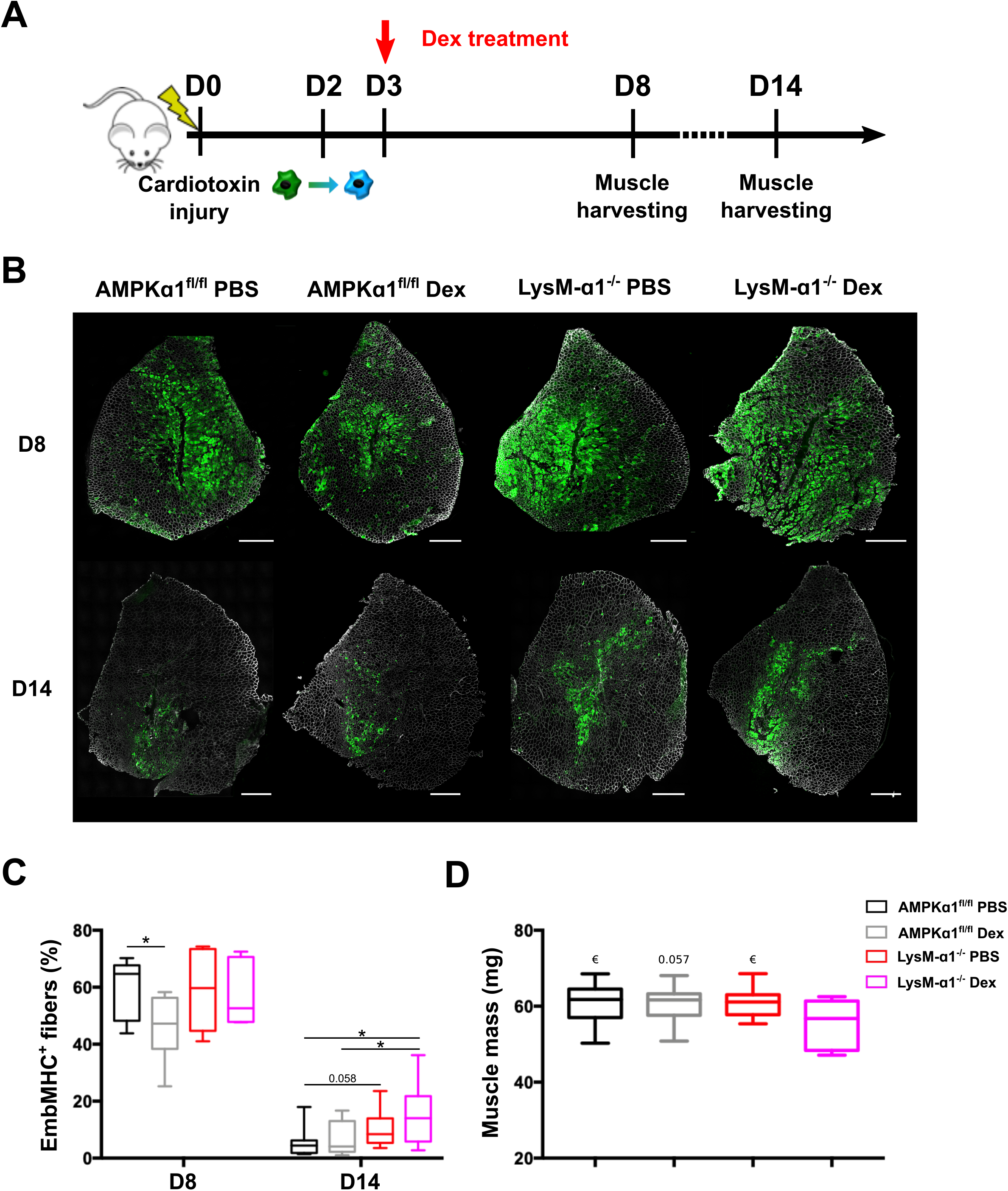
AMPKa1 in macrophages is required for glucocorticoid-induced effect on maturation of regenerating myofibers. **(A)** Experimental procedure of mice treatment with Dex. After cardiotoxin injection to damage the muscle, AMPKα1^fl/fl^ (WT) and LysM-α1^−/−^ mice were treated with a single dose of dexamethasone (Dex) intra-peritoneal (i.p.) (0.1 mg/kg) at day 3 (D3) and *Tibialis anterior (TA)* muscles were harvested at day 8 (D8) and 14 (D14) after injury. **(B)** Representative embryonic myosin heavy chain (EmbMHC) staining on whole *TA* muscle section at D8 and D14 after injury. **(C)** Quantification of **(B)**, results are expressed as percentage of positive myofibers expressing the EmbMHC on the whole muscle section. **(D)** Muscle mass at day 14 after cardiotoxin injury. Results are means ± SEMs of 7-12 muscles (C-D, D8 & D14). * p<0.05, € p<0.05 *vs* LysM-α1^−/−^ Dex using Kruskal Wallis (C) or ANOVA (D) tests. Bar= 500 μm (B).

### AMPKα1 mediates glucocorticoid induced macrophage efferocytosis and shift toward a restorative inflammatory profile

Next, we aimed to understand how AMPK loss influences the modulation of inflammatory activated macrophages in the two different models of inflammation. To mimic infectious inflammatory condition, we treated BMDMs with either LPS or with damage-associated molecular patterns (DAMPs, present in early damaged skeletal muscle) to mimic sterile inflammatory conditions (i.e. tissue damage conditions) for 24 hours and we added Dex to the cells for 6 hours before analysis. We confirmed that BMDMs treated for 24 hours with muscle DAMPs acquired a pro-inflammatory phenotype, assessed by the increased expression of the pro-inflammatory markers iNOS and CCL3 and the downregulation of the anti-inflammatory marker CD206 (Fig.S4A). We then compared genes differentially expressed between WT and AMPKα1^−/−^ BMDMs in DAMPs+Dex and LPS+Dex conditions by RNA-seq. We identified an overlap of 76 genes, that were commonly up-regulated by LPS+Dex and DAMPs+Dex in WT but downregulated in AMPK-deficient macrophages. This molecular signature suggests a function of AMPK and GR signaling across TLR4, LPS induced inflammatory macrophage pathways, and the DAMPs induced macrophage inflammatory pathways (Fig.4A, Supplementary Table 2). Among these down-regulated genes, 57 were annotated and Gene Ontology analysis identified pathways that regulate the inflammatory status/polarization of macrophages, such as arginine metabolism, oxidative phosphorylation, PPARγ signaling and the HIF1α signaling pathway (Cramer et al., 2003; Varga et al., 2016b) (Supplementary Table 2). We also found overlap in expression of *Klf4*, *Nr1h3* and *Il10rb* genes, that are directly involved in anti-inflammatory polarization (Kourtzelis et al., 2019; Liao et al., 2011; Shouval et al., 2014). Among the 57 genes having a known function, 22 were involved in either phagocytosis, autophagy and/or macrophage polarization (Supplementary Table 2). Phagocytosis of apoptotic cells, also known as efferocytosis, is a core function of macrophages during tissue repair and inflammation, and has been shown to be a key factor in the resolution of acute lung inflammation (Fadok et al., 1998; Nepal et al., 2019), and for efficient tissue repair (Koroskenyi et al., 2011).

**Figure 4.**
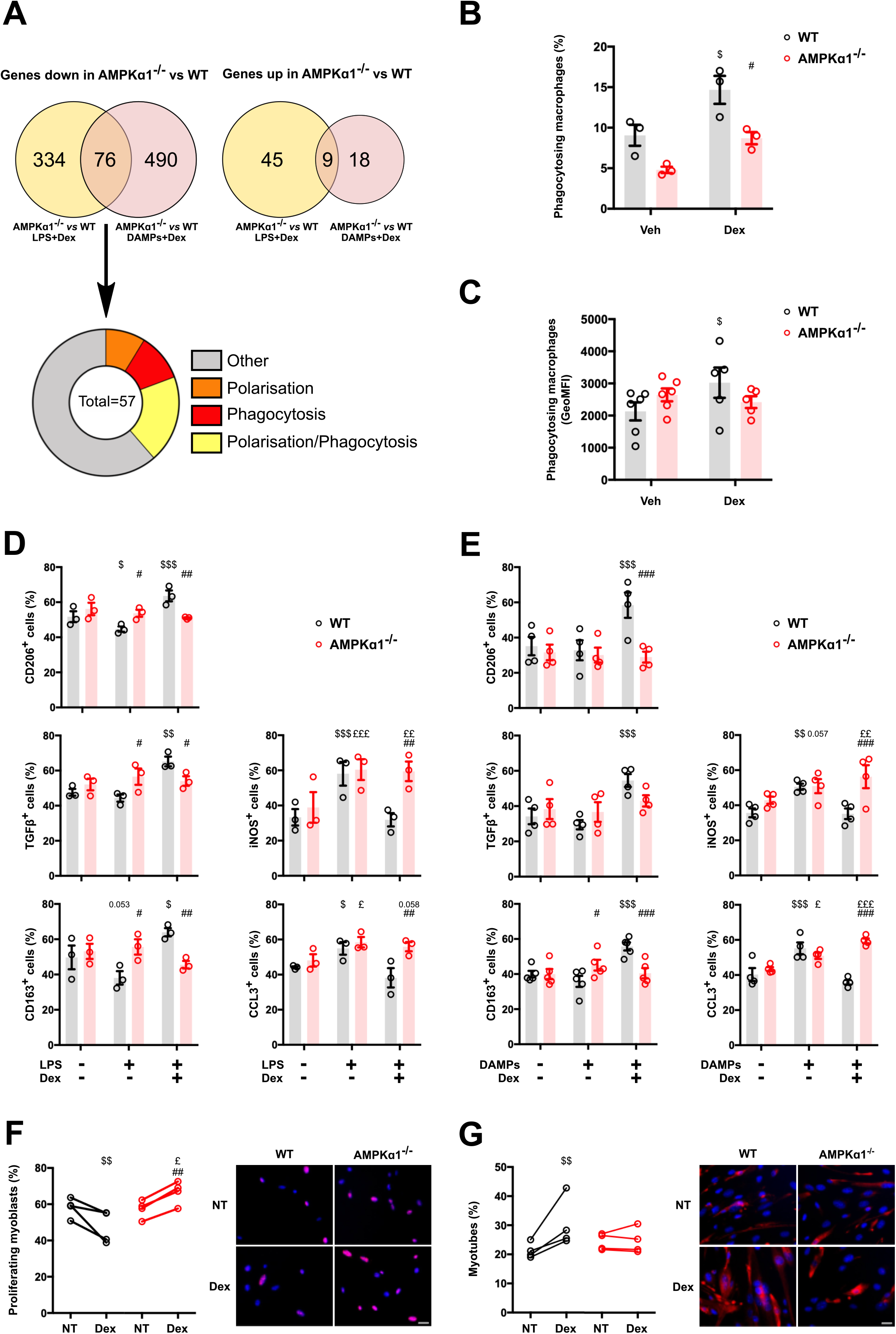
AMPKa1 mediates glucocorticoid-induced macrophage acquisition of an anti-inflammatory phenotype. **(A)** Venn diagrams from RNAseq experiments indicating genes differentially expressed in AMPKα1^−/−^ *vs* WT macrophages treated with either LPS or Damaged Associated Molecular Patterns (DAMPs) for 24 hours and further treated with dexamethasone (Dex). Genes downregulated in AMPK deficient macrophages were analyzed for their functions using Gene Ontology. **(B)** Bone marrow-derived macrophages (BMDMs) were treated or not with Dex and cultured with CSFE-fluorescently labelled apoptotic thymocytes for 2.5 hours. The number of macrophages having ingested apoptotic cells was evaluated by flow cytometry. **(C)** *Tibialis anterior* (TA) muscle from WT and AMPKα1^−/−^ mice was damaged with cardiotoxin and Dex (or vehicle) was *i.p.* injected around 60 h after injury (~ 2.5 days) prior CD45^pos^ cells were sorted with magnetic beads 15 hours later (day 3). CD45^pos^ CD64^pos^ Ly6C^pos^, Ly6C^int^ and Ly6C^neg^ macrophage subsets were cultured with dead fluorescently labelled myoblasts for 6 hours. Phagocytosis was evaluated by flow cytometry and expressed as the GeoMean fluorescence Intensity of the phagocytosing cells (see FigS4B). **(D-E)** WT and AMPKα1^−/−^ BMDMs were treated with either LPS **(D)** or muscle DAMPs **(E)** for 24 hours and further treated with Dex for 6 hours before immunofluorescence for anti- (CD206, TGFβ, CD163) and pro-inflammatory (iNOS, CCL3) markers was performed. Results are expressed as percentages of positive cells. **(F-G)** WT and AMPKα1^−/−^ BMDMs were treated with Dex for 72 hours, washed and the conditioned medium was recovered after 24 hours and tested on muscle stem cells that were tested for their proliferation using Ki67 immunolabeling **(F)** or their differentiation using desmin labeling and counting of the number of myonuclei in the cells **(G)**. Results are means ± SEMs of 5-6 (C), 3 (B,D) or 4 (E-G) experiments. $ p<0.05, $$ p<0.01, $$$ p<0.001 *vs* untreated WT; £ p<0.05, ££ p<0.01, £££ p<0.001 *vs* untreated AMPKα1^−/−^; # p<0.05, ## p<0.01, ### p<0.001 compared AMPKα1^−/−^ *vs* WT for each condition, using or ANOVA (B-G). Bar=10 μm (F,G).

To determine whether AMPK alters the ability for GCs to induce efferocytosis, we performed *in vitro* and *ex vivo* efferocytosis assays. While Dex treatment increased efferocytosis of apoptotic thymocytes (simulating cells recruited during infections such as sepsis) in WT BMDMs (+62%), AMPKα1 deficient macrophages were unable to efficiently phagocytose the apoptotic cells in response to Dex stimulation (Fig.4B). This was confirmed *ex vivo*, in a sterile inflammation model where CD64^pos^ macrophages were analysed from post-injured regenerating skeletal muscle and were tested for their capacity to engulf apoptotic myoblasts after a single Dex injection during the resolution of inflammation, *i.e*. at 2.5 days (Mounier et al., 2013). Three subsets of macrophages were sorted as: pro-inflammatory Ly6C^pos^ macrophages, Ly6C^neg^ anti-inflammatory macrophages and a Ly6C^int^ subset of macrophages that are shifting from pro- to anti-inflammatory profile (Mounier et al., 2013; Varga et al., 2013; Varga et al., 2016a; Varga et al., 2016b) (Fig.S4B). In WT mice, Dex treatment of mice 15 hours prior to macrophage isolation increased by +42% the phagocytic capacity of Ly6C^int^ macrophages assessed by the increased GeoMean Fluorescent Intensity of macrophages having engulfed apoptotic myoblasts (Fig.4C, Fig.S4B). On the contrary, in AMPKα1 deficient animals, Ly6C^int^ macrophages from regenerating muscle did not modify their phagocytic capacity upon Dex treatment (Fig.4C, Fig.S4B). Of note, Ly6C^pos^ and Ly6C^neg^ did not show changes in their phagocytic capacity upon the various conditions (Fig.S4B). These data are in accordance with the impaired gene regulation in AMPK deficient macrophages upon Dex treatment and indicate that loss of AMPK abrogates GC-induced efferocytosis, a major process contributing to the resolution of inflammation.

Next, we tested whether the AMPK-dependent increased capacity of efferocytosis was associated with an acquisition of a pro-resolving inflammatory profile upon Dex treatment. WT and AMPKα1^−/−^ BMDMs were treated with LPS (Fig.4D) or with DAMPs (Fig.2E) for 24 hours, then Dex was added for 6h and immunofluorescence for pro- and anti-inflammatory macrophage markers was performed. WT BMDMs responded to Dex treatment by increasing the expression of the anti-inflammatory markers CD206, TGFβ and CD163 (+43, +47, +68% in LPS+Dex *vs*. LPS treated cells and +78, +85, +55% in DAMPs+Dex *vs*. DAMPs treated cells, respectively) and decreasing that of the pro-inflammatory markers iNOS and CCL3 (−45,−30% in LPS+Dex *vs*. LPS treated cells and −30, −35% in DAMPs+Dex *vs*. DAMPs treated cells, respectively) (Fig.4D,E). On the contrary, LPS- or DAMPs-triggered pro-inflammatory AMPKα1^−/−^ BMDMs did not shift their inflammatory profile upon Dex treatment (Fig.4D,E). These results indicate that AMPK is required for the acquisition of the anti-inflammatory profile induced by GCs. We then further analyzed whether this phenotype is associated with macrophage function, using conditioned medium from Dex-treated WT and AMPKα1^−/−^ BMDMs applied to muscle cells (i.e. myoblasts). We previously showed that this *in vitro* model is a reliable read out to compare the function of pro- and anti-inflammatory macrophages that stimulate myoblast proliferation and differentiation, respectively (Mounier et al., 2013; Varga et al., 2016b). Medium from Dex-treated WT macrophages decreased myoblast proliferation (−20%) (Fig.4F) and increased myoblast differentiation (+37%) (Fig.4G) indicating a shift toward a pro-resolving phenotype, as expected. However, these properties were lost in AMPKα1^−/−^ macrophages, either at 10^−7^M or 10^−8^M (Fig.4F,G, Fig.S4C,D). Altogether, these data demonstrate that loss of AMPKα1 alters GC regulated transcriptional activity in macrophages associated with efferocytosis, diminishing the potential for GR to regulate the acquisition of a functional anti-inflammatory phenotype through the alteration of the efferocytic properties of macrophages.

### An AMPK-FOXO3-GR axis regulates glucocorticoid mediated gene expression, and macrophage polarization

In order to decipher the molecular mechanism underlying this regulation, we searched for transcription factor binding sites enriched in genes differentially regulated by Dex in WT and in AMPKα1^−/−^ BMDMs. As expected, GR consensus binding site was enriched in genes commonly activated by Dex in WT and AMPKα1^−/−^ macrophages, validating the relevance of this strategy. Interestingly, binding sites for the transcription factors *Sry*, *Nkx2-5*, *Foxd1* and *Foxo3* were specifically enriched in genes up-regulated by Dex only in WT cells (Fig.S5) together with genes down-regulated by Dex only in AMPKα1^−/−^ cells (Fig.S6). This suggested a differential role for these transcription factors in Dex-mediated transcriptional regulation in the presence or absence of AMPKα1. Of note, within transcription factors known to be involved in macrophage function (FOXO3, HIF1:ARNT, PPARγ:RXRα, GR and AR), only FOXO3 followed a clear pattern of enrichment found in common and WT-selective up-regulated genes, and AMPKα1^−/−^ selective down-regulated genes (Fig.5A). To further confirm the involvement of FOXO3 in AMPK-dependent GR responses, we used Gene Set Enrichment Analysis using dex-regulated genes in WT and AMPKα1^−/−^ macrophages, and a gene set of FOXO3-dependent genes defined by FOXO3 KO macrophages (Litvak et al., 2012) (Fig.5B). We found that WT cells had a significant enrichment for FOXO3-dependent genes, while dex regulated genes in AMPKα1^−/−^ macrophages did not. These data suggest a potential co-regulatory role of FOXO3 and GR, requiring AMPK for gene expression. Therefore, we assessed whether FOXO3 DNA loading was GC regulated, in both WT and AMPK deficient cells by ChIP-PCR, at dex regulated genes that were upregulated only in WT macrophages (Fig.5C). We used publicly available ChIP-seq data to choose sites where GR binds to determine whether FOXO3 binds at similar loci (Fig.S7). Strikingly, FOXO3 was recruited to similar loci as GR in a GC-dependent manner, however this effect was abolished in the AMPK deficient cells (Fig.5C), further establishing the link between AMPK, GR and FOXO3. We next determined whether activation of FOXO3 could recover the detrimental effect of AMPK deletion on GC activity in pro-inflammatory macrophages. As there are no known FOXO3 agonists, we took advantage of the AKT inhibitor MK2206, and activated FOXO3 by inhibiting its inhibitor AKT. Indeed, AKT phosphorylates, and sequesters FOXO3 in the cytoplasm, preventing FOXO3 transcriptional functions (Brunet et al., 1999). We first treated WT and AMPKα1^−/−^ BMDMs with DAMPs or LPS for 24 h and added either vehicle or MK2206 for 1 hour before further treatment with dex for 6 h. We then assessed the expression of the pro- and anti-inflammatory markers iNOS and CD206 (Fig.5D, E). The combinatorial treatment of MK2206 and dex prevented the deleterious effect of AMPKα1 deletion after inflammatory stimulus on the expression of iNOS. Similarly, CD206 expression was also recovered, with MK2206 treated AMPKα1^−/−^ cells expressing significantly more than their untreated counterparts. This result matched the CD206 levels of DAMPs+dex or LPS+dex WT cells. These data demonstrate that indirect FOXO3 activation through AKT inhibition can overcome the loss of AMPK in macrophages, suggesting a signaling pathway consisting of AMPK, GR and FOXO3, to regulate macrophage polarization (Fig.5F).

**Figure 5.**
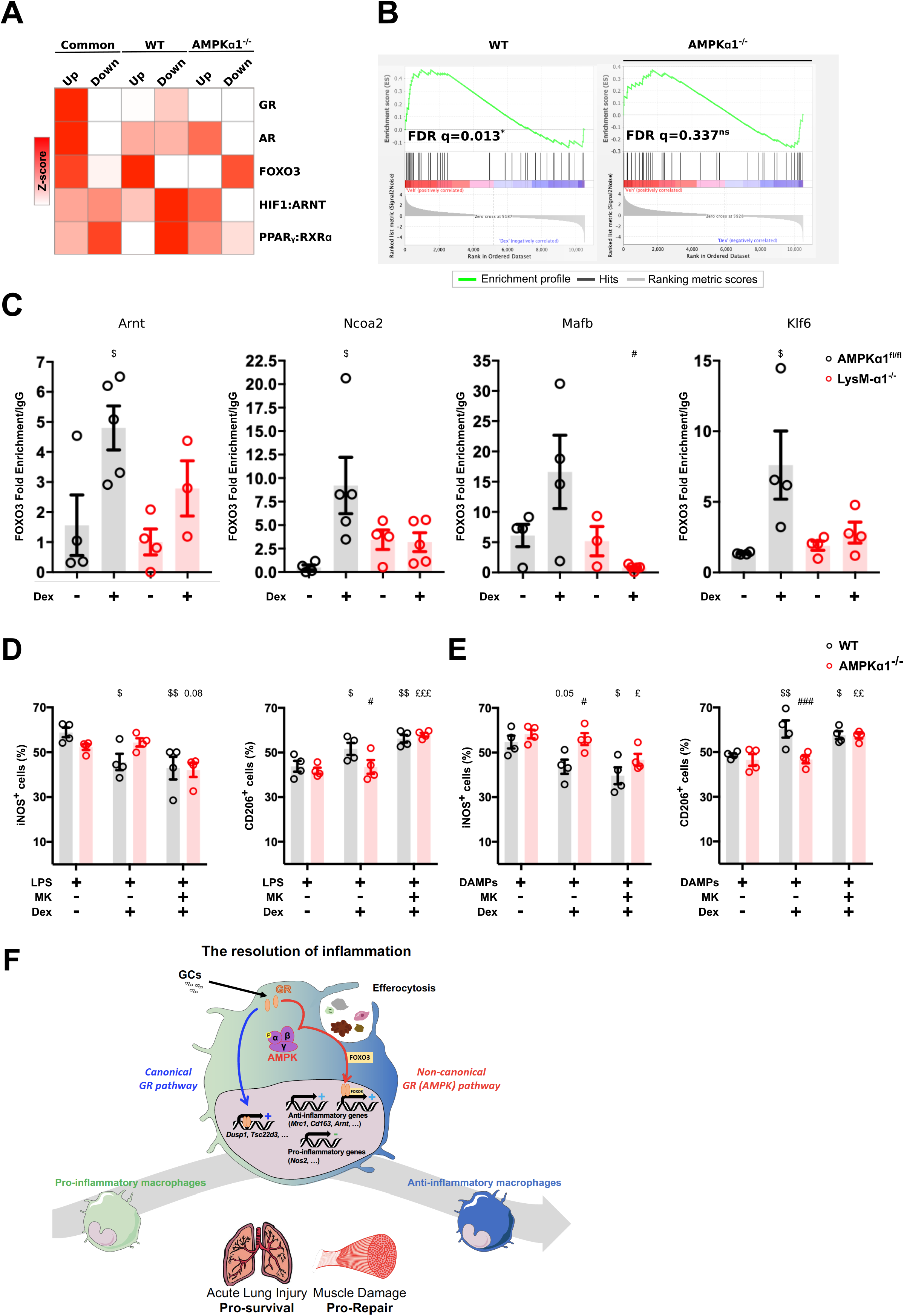
An AMPK-GR-FOXO3 Trinity Regulates Glucocorticoid Induction of Anti-inflammatory Macrophages. **(A)** Transcriptional regulators of differentially regulated identified by oPOSSUM were stratified into known regulators of macrophage function. **(B)** GSEA analysis of dexamethasone (Dex) regulated genes in WT and AMPKα1^−/−^ Bone marrow-derived macrophages (BMDMs) for FOXO3 dependent targets. **(C)** ChIP-PCR analysis of FOXO3 binding in AMPKα1^fl/fl^ (WT) and LysM-α1^−/−^ BMDMs. **(D-E)** WT and AMPKα1^−/−^ BMDMs were first treated with either LPS **(D)** or muscle Damaged Associated Molecular Patterns (DAMPs) **(E)** for 24 hours. Then, cells were treated with MK2206 for 1 hour prior addition of Dex for 6 hours before immunofluorescence for anti- (CD206) and pro-inflammatory (iNOS) markers was performed. (**F**) Graphical representation of how signaling pathway AMPK/GR/FOXO3 axis could regulate macrophage polarization during the resolution of inflammation. Results are expressed as percentages of positive cells. Results are means ± SEMs. $ p<0.05, $$ p<0.001 *vs* control WT, £ p<0.05, ££ p<0.01, £££ p<0.001 *vs* control AMPKα1^−/−^, # p<0.05, ### p<0.001 compared AMPKα1^−/−^ *vs* WT, using Mann-Whitney-Wilcoxon test (A) or ANOVA and Kruskal-Wallis tests (C-D-E).

## Discussion

Here we demonstrate that GCs require AMPKα1 to efficiently polarize macrophages to an anti-inflammatory phenotype. GCs were able to activate AMPK in macrophages, and genetic deletion of AMPKα1 inhibited GR mediated anti-inflammatory macrophage polarization and functions. Intriguingly, regulation of well-known GR target genes were not affected, whereas a novel subset of genes involved in polarization and efferocytosis required AMPK activation. This had important implications: i) *in vitro,* where GC-induced phagocytosis was abrogated in the absence of AMPK leading to defective acquisition of the anti-inflammatory status phenotype and functions, and ii) *in vivo* where GC treatment manifested in a delayed onset of skeletal muscle regeneration and inefficient inhibition of lung inflammation in LysM-α1^−/−^ mice, suggesting a key role of AMPK in mediating the anti-inflammatory effects of GCs in macrophages.

Only scant studies have reported a link between AMPK signaling and GR signaling, and all in metabolic organs and cells (Liu et al., 2016; Nader et al., 2010; Ratman et al., 2016). Indirect AMPK activation is observed in muscle cells where dexamethasone treatment induces mitochondrial dysfunction leading to decreased ATP, thus activating AMPK (Liu et al., 2016). AMPK indirectly phosphorylates GR S211 *via* p38 in preadipocytes and hepatocytes, promoting GR transcriptional activity (Nader et al., 2010). However, our results show that in macrophages, GR phosphorylation depended on AMPK completely separate from the p38 pathway. In addition, dexamethasone was shown to increase AMPK activity in hepatocytes while decreasing it in adipocytes (Christ-Crain et al., 2008). Finally, a physical link between phospho-AMPK and phospho-GR was shown in complex with PPARα to increase the transcriptional activity of the latter upon starvation (Ratman et al., 2016). These studies show that AMPK-GR interaction impacts on GR transcriptional activity. In our conditions, we found that GCs activate AMPK as efficiently as the potent and allosteric AMPK activator 991 in macrophages, indicating a previously unknown role of GCs in stimulating AMPK signaling.

Deletion of AMPK results in a profound change of Dex-mediated gene expression. Interestingly, canonical and well-studied GR related genes (both up and down regulated) were not affected by the loss of AMPK in macrophages. The induction of GC response element (GRE) containing genes such as *Gilz* (*Tsc22d3*), *Mkp1* (*Dusp1*) and *Fkbp5* or the repression of genes with elements required for tethering such as *Tnfa, Il6* and *Ifng*, were not affected by the loss of AMPK in macrophages and in the BAL during lung inflammation. Importantly, we did find a difference in the capacity for GC signaling in absence of AMPK to polarize macrophages towards an anti-inflammatory phenotype. For example, we found key genes, such as *Klf4, Nr1h3* and *Il10rb* all down-regulated in the AMPK knockout macrophages after treatment with LPS or DAMPs and Dex. We also saw an effect of GCs on the expression of *Lamtor* subunits 1-5, down regulating them in an AMPK dependent manner. The LAMTOR complex is a known regulator of mTOR, with genetic ablation of LAMTOR decreasing mTOR activity in immune cells (Hosokawa et al., 2017). This effect was only observed in WT cells, indicating a cross-talk between GCs and AMPK in regulating mTOR, known to be involved in promoting pro-inflammatory macrophage polarization (Byles et al., 2013). Similarly, GC mediated upregulation of the GR co-activators *Ncoa2, Ncoa5* and *Ncoa6* was dependent on intact AMPK signaling. NCOA proteins are known to control the target and directionality of gene regulation by GR (Petta et al., 2016), thus as AMPK regulates the expression of these, adding a further layer of AMPK-mediated regulation of GC signaling. Interestingly, we found also a substantial fraction of genes that were Dex down-regulated in the absence of AMPK, but not in WT cells. This Dex regulation unleashed in AMPK knockout macrophages was affecting genes annotated for regulation of proliferation and mitosis. This could be interesting for macrophage homeostasis since restorative/anti-inflammatory macrophages were shown to proliferate at the time of the resolution of inflammation during tissue repair (Arnold et al., 2007). Since we focused in this study on pro- *versus* anti-inflammatory functions of macrophages, we did not follow this up in this study. GR phosphorylation at S211 may also be involved in the AMPK control of non-canonical GR signaling. Previous reports have shown that phospho-deficient (S211A) and phospho-mimetic (S221D) have opposite effects on GR transcriptional activity, and on binding to the MED14 subunit of the Mediator complex (Chen et al., 2008). While the authors only confirmed one interaction, the implication remains that differential phosphorylation status of GR, regulated by different kinases, can alter interaction with other co-factors, and thus the target genes of GR (Galliher-Beckley and Cidlowski, 2009). Indeed, this also applies to the phosphorylation status of GR co-factors, for example GRIP-1 phosphorylation can alter the transcriptional activity of GR (Rollins et al., 2017). However, the question of how AMPK can interact with various GR cofactors is still open.

While the above-mentioned genes illustrate an AMPK-dependent regulation of non-canonical GR transcriptional activity, our analysis of the enrichment of genes exhibiting specific transcription factor binding sites indicated that FOXO3 binding site was found in genes upregulated in WT macrophages upon Dex treatment and in genes downregulated in AMPK KO macrophages. FOXO3 is a direct target of AMPK (Greer et al., 2007; Lutzner et al., 2012) proposed that upon metabolic stress (high cortisol and high AMP), FOXO3 is an indirect GR target through the activation of SGK1 that alleviates the inhibition of FOXO3 transcriptional activity, which is synergized with the activation of FOXO3 transcriptional activity by AMPK. FOXO3 was shown to be required for the differentiation of monocytes into macrophages *via* activation of AMPKα1 and autophagy (Zhang et al., 2017) and to promote anti-viral response in macrophages (Zhang et al., 2019b). Strikingly, FOXO3 is involved in the GC-driven anti-inflammatory response in Systemic Lupus Erythematosus through the reinforcement of the blockage of NF-KB activity (Lu et al., 2016). Indeed, GR and FOXO1, a related Forehead box transcription factor, are found at similar loci in the liver, suggesting that GR-FOXO dependent gene regulation occurs across multiple transcription factors in different organs (Kalvisa et al., 2018). These observations open up the possibility that either GR or FOXO3 are assisting one-another in binding the DNA and regulating macrophage alternative activation genes, using an assisted loading mechanism. We did not define whether GR or FOXO3 is acting as the pioneer factor, partly due to the complex molecular links between AMPK activation by GCs, and AMPK-dependent activation of FOXO3. It is likely that GCs, through AMPK, promote FOXO3 nuclear accumulation, along with GR nuclear translocation simultaneously. However, the clear implication of an AMPK-GR-FOXO3 regulatory trinity, is that feeding/fasting can control macrophage functionality. Through multiple hormonal signals such as cortisol and insulin, which activate GR and inhibit FOXO3 respectively, along with changes in cellular AMP levels, inflammation can be modulated by endocrine control of organismal metabolism. Supporting the notion that FOXOs are important for resolution of inflammation, insulin treatment has been shown to exacerbate sepsis-induced lung injury in diabetic rats (Filgueiras et al., 2014). Moreover, previous reports have shown that GCs control skeletal muscle metabolism, through the induction of the transcription factor KLF15, which activity was a key component for the repair in a mouse model of Duchenne muscular dystrophy (Morrison-Nozik et al., 2015). The metabolic actions of GCs may therefore be highly unappreciated in the regulation of inflammatory diseases, but our data further support the necessity for metabolic modulation as a core aspect of GCs in the resolution of inflammation.

GC/GR signaling enhances the efferocytic properties of macrophages (Giles et al., 2001), for instance, by increasing the expression of *Mertk* or *Anxa1* as a core aspect of their anti-inflammatory effects (Koenen et al., 2018; Zizzo et al., 2012). Efferocytosis is associated with macrophage polarization state since it induces the acquisition of an anti-inflammatory/resolving phenotype (Fadok et al., 1998; Zhang et al., 2019a), thus provides a bridge in macrophage function across sterile and LPS induced inflammation. In that context, our results show that genes downregulated in AMPK-deficient macrophages upon GC treatment were associated with autophagy and/or phagocytic processes. We previously showed that AMPK is required for the shift of macrophage phenotype upon phagocytosis of apoptotic cells (Mounier et al., 2013). Here we show that functionally, AMPK-deficient macrophages did not increase their phagocytic activity upon GC treatment. Consequently, LPS- or DAMPs-treated AMPK-deficient macrophages did not respond to GCs and were not able to shift towards an anti-inflammatory profile nor to acquire the promyogenic function of anti-inflammatory macrophages.

*In vivo*, the consequence of an impaired acquisition of an inflammatory/resolving phenotype of macrophages upon GC treatment results in a delay and alteration of the return to homeostasis in a model of tissue repair, assessed by the alteration of myosin heavy chain expression in regenerating skeletal muscle. It also strongly impacts the response of the tissue to a severe acute inflammation (in a lung injury model) where AMPK is required for macrophages to respond to Dex treatment in order to dampen the inflammation and recover the tissue integrity, and survival. These results implicate a convergence, and interdependence of GR and AMPK in the anti-inflammatory response that is independent on reduction of cytokine expression and is therefore a so far overlooked important aspect of mechanism of GC usage in the clinics.

Altogether, our data demonstrate a dependence of GR on AMPK and FOXO3 in the regulation of inflammation, through a non-classical pathway in macrophages. This has clear implications for the role of GR signaling on macrophage function depending on the energy status of the cell, and could help identification of new small molecule strategies to enhance GC action in a clinical setting.

## Acknowledgments

This work was supported by French-German Collaboration for Joint Projects of the French Agence Nationale de Recherce (ANR) and the Deutsche Forschungsgemeinschaft (DFG) to BC and JT (Tu220/13), Campus France-DAAD to BC and JT, CNRS, Société Française de Myologie and by Fondation pour la Recherche Médicale (Equipe FRM DEQ20140329495), JT was supported by CRC1149 Trauma and GC supported by Bausteingrant from the Medical Faculty of Ulm University and ProtrainU grant from Ulm University. Excellent support was supplied by the animal facilities at University of Ulm and the Université Claude Bernard Lyon 1.

## Experimental procedures

### Animal models

C57BL/6, LysM^cre/+^;AMPKa1^fl/fl^ (B6-Prkaa1^flox/flox^/J; B6-Lyz2^tm1(cre)ifo^, designated as LysM-α1^−/−^) and 129;B6-Prkaa1^tm^ (AMPKα1^−/−^) adult male mice (8 to 14-weeks-old) were bred and used according to French and German legislations and protocols were approved by ethical committees. For muscle injury experiments, mice were anesthetized with isoflurane and *Tibialis Anterior* (TA) muscle was injected with cardiotoxin (Latoxan) at 12 μM (50 μl *per* muscle) as previously described (Mounier et al., 2013). Mice were treated by intra-peritoneal (i.p.) injection of either dexamethasone (Dex) (concentrations indicated in the figures, Sigma-Aldrich, D2915) or PBS (Vehicle) at various time points after injury. Muscles were harvested at different time points after injury. For lung inflammation experiments, male and female mice (8-14 weeks) were injected i.p. with either vehicle (PBS) or LPS (Sigma-Aldrich, L2880) at a concentration of 10 mg/kg. Mice were then injected with either Dex (Sigma-Aldrich, D2915) 1mg/kg or cyclodextrin (1 mg/kg) vehicle control. After 30 minutes, mice were injected intravenous (i.v.) with 2.6 μl/g oleic acid (Sigma-Aldrich, O1008) as a solution of 40% oleic acid in 1% bovine serum albumin (Sigma-Aldrich, A3912) as previously described (Vettorazzi et al., 2015). Mice were killed 24 h later and bronchoalveolar lavage (BAL) was performed by instilling the lungs with 1mL of BAL fluid (PBS, 10 mM EDTA, 1% BSA) 3 times. Survival after lung inflammation was measured for 53 h. Mice body temperature and body mass were measured each 4 h during 48 h after induction of lung injury. Mice were sacrificed when limit points were reached, body temperature lower than 29°C and 15 % of total body weight loss. General aspect of mice was also considered.

### Analysis of lung immune cells, total cell counts and FACS

Cells isolated by BAL were centrifuged, washed with BAL fluid and red blood cells lysed. Resulting cells were counted using a hemocytometer and normalized to the total amount of BAL fluid recovered. For FACS, cells were suspended in BAL fluid containing FcɣRII/III block (eBiosciences, 1:300), washed then incubated with antibodies against CD45 (APC, Invitrogen 30-F11, 1:300), CD11b (FITC, Invitrogen M1/70, 1:300), F4/80 (AF700, Invitrogen BM8, 1:300), Ly6G (Pe-Cy7, Invitrogen RB6-8C5, 1:300), Ly6C (PerCP-Cy5.5, eBiosciences HK1.4, 1:300). After washing, cells were analyzed immediately using a flow cytometer (LSRII, BD). Data were analyzed using FlowJo software.

### Cytokine multiplex

Cytokines were determined in BAL fluid using a Cytokine & Chemokine Convenience 36-Plex Mouse ProcartaPlex Panel 1A (ThermoFisher) using a Bio-plex 200 System (BioRad). Samples below the detection threshold were excluded from analysis.

### Histological analysis and *in vivo* immunostaining

TA muscles were mounted on piece of cork and fixed with tragacanth gum (VWR, 24437.260) before being frozen in isopentane cooled by liquid nitrogen. Muscles were stored at −80°C. TA muscles were sectioned with cryomicrotome (NX50, Microm) in 10 μm thick sections and slides were kept at −80°C. Slides were first stain with Hematoxilin and Eosin to check cardiotoxin injection and immunofluorescence was performed only on muscles that showed at least 90% of their area occupied by regenerating myofibers (*i.e.* exhibiting central nuclei). For immunofluorescence experiments, slides were incubated with Triton x100 0.5%, saturated with bovine serum albumin (BSA) 2% and were incubated overnight with primary anti-body: anti-embMHC, (Santa Cruz, sc-53091, 1:100) or anti-laminin (Sigma Aldrich, L9393, 1:200). Slides were washed with PBS and incubated with secondary antibodies: Alexa Fluor 488 anti-mouse IgG1 (Jackson Immuno research, 115-545-205, 1:200) or Cy5 Donkey anti-Rabbit IgG (H+L) (Jackson Immuno research, 711-175-152, 1:200). Sections were soaked for 10 s in Hoechst solution H 33342 (Sigma-Aldrich, B2261, 1:1000) and washed once with PBS before mounting with antifading Fluoromount G medium (Interchim, FP-483331). Slides were stored at 4°C protected from light until picture acquisition. For adult myosin immunostaining, triton was not used and slides were blocked with Mouse on Mouse (M.O.M™) Blocking Reagent (Vector Laboratories, MKB-2213). All samples of the same experiment were recorded in similar conditions (microscope, magnification, exposure time and binning). For whole section analysis, slides were automatically scanned at ×10 or ×20 of magnification using an Axio Observer.Z1 (Zeiss) connected to a CoolSNAP HQ2 CCD Camera (Photometrics) using Metavue software. The whole muscle section was automatically reconstituted by Metavue software. Cross sectional analysis was performed using Open-CSAM ImageJ macro (Desgeorges et al., 2019b).

After BAL, lungs were fixed in 4% PFA overnight, then transferred to 70% ethanol. Fixed lungs were processed and embedded into paraffin, then cut to 5 μm sections using a microtome (Leica RM2255). Slides were then stained with hematoxylin and eosin, and imaged using an Olympus BX41 microscope at 20X magnification.

### Bone marrow-derived macrophages

Macrophages were obtained from bone marrow precursors as previously described (Mounier et al., 2013). Total bone marrow was isolated from mice by flushing the femur and tibiae bone marrow with culture medium. Extracted bone marrow was cultured in DMEM high glucose high pyruvate (Gibco, 11995065) containing 20% of FBS, 1% of penicillin/streptomycin, 1% of fungizone and 30% of L929 conditioned medium for 5 to 7 days. L929 secrete high levels of Macrophage Colony-Stimulating Factor (MCSF) that triggers macrophage differentiation. L929 conditioned medium was obtained after 10 days of L929 cells culture with DMEM containing 10% FBS.

### DAMPs – Damaged associated molecular patterns from muscle

DAMPs were obtained from ischemia-injured muscles. Briefly, mice hindlimb underwent an ischemia for 2 hours. After 20 hours of reperfusion, all hindlimb muscles were harvested. Three mice were pooled for each substrate. Once harvested, muscles were digested with Mammalian cell lysis buffer (Sigma-Aldrich, MCL1). Two tungsten beads were added in samples to homogenize the muscle extract with a Tissue lyser. After centrifugation, samples were sonicated and supernatant was recovered, pooled, and stored at −80°C.

### Macrophage inflammatory status

Macrophages were seeded at 54,000 cells/cm^2^ in DMEM containing 10% of FBS on glass coverslips in 12-well plates. Cells were treated with muscle DAMPs (1 μg/ml) or LPS (100 ng/ml for 24 hours and Dex (Sigma-Aldrich, D4902, 1×10^−8^ M) was further directly added in the well for 6 hours. In the AKT inhibition experiments, after the 24 hour incubation with muscle DAMPs or LPS, cells were treated with MK-2206 (Adooq Bioscience, A10003, 1 μM) (or 0.01% of DMSO for controls) for 1 hour prior addition of Dex.

Cells were fixed with PFA 4%, permeabilized with Triton x100 0.5% and incubated for 1 hour with BSA 4%. Staining for CCL3 (Santa Cruz, sc-1383, 1:50), iNOS (Abcam, ab3523, 1:50), CD206 (Santa Cruz, sc-58987, 1:50), CD163 (Santa Cruz, sc-33560, 1:50 or BIOSS, bs-2527R, 1:100), TGFβ1 (Abcam, ab64715, 1:50) was performed as in (Mounier et al., 2013). All samples of the same experiment were recorded in similar conditions (microscope, magnification, exposure time and binning). At least 10 pictures *per* condition were acquired at x20 magnification on an Axio Imager.Z1 microscope (Zeiss) connected to a CoolSNAP MYO CCD camera (photometrics) using MetaMorph Software. Quantification of macrophage markers was done manually using ImageJ software.

### Quantitative Real-Time PCR and RNA-sequencing

Macrophages were seeded and allowed to adhere for 16 hours. Cells were then washed twice with PBS and medium was changed to Macrophage-SFM (Thermofisher, 12065074) with 1% penicillin/streptomycin, 1% L-glutamine, 1% sodium pyruvate, 1% fungizone for 16 hours. Cells were then treated with either vehicle (DMSO), 100 ng/mL LPS (Sigma, L6529), or DAMPS (1 μg/ml), 100 nM Dex (Sigma, D4902), or a combination of both for either 24, or 6 hours. Cells were then washed twice with PBS and total RNA extracted using Trizol (Invitrogen, 15596018), according to the manufacturer’s protocol. For qRT-PCR, RNA was measured using a Nanodrop 1 μg of RNA was then reverse transcribed into cDNA (Applied Biosystems, High Capacity cDNA kit, 4368813) with additional RNase inhibitor (Invitrogen RNase OUT, 10777019). cDNA was quantified using a ViiA 7 (Applied Biosystems) and analyzed using the delta-delta CT method, relative to the average of WT vehicle treated samples. For RNA-seq, RNA integrity was assessed using a 2100 Bioanalyzer (Agilent) and samples with a RIN value of >6 were taken forward for library prep. Library prep and sequencing was performed using the Novaseq 6000 S4 platform at Novogene (Novogene, Beijing, China). Differential expression analysis was performed using DESeq2. Gene ontology and pathway analyses were performed using DAVID (Huang da et al., 2009), and ENRICHR (Kuleshov et al., 2016).

### Primer List

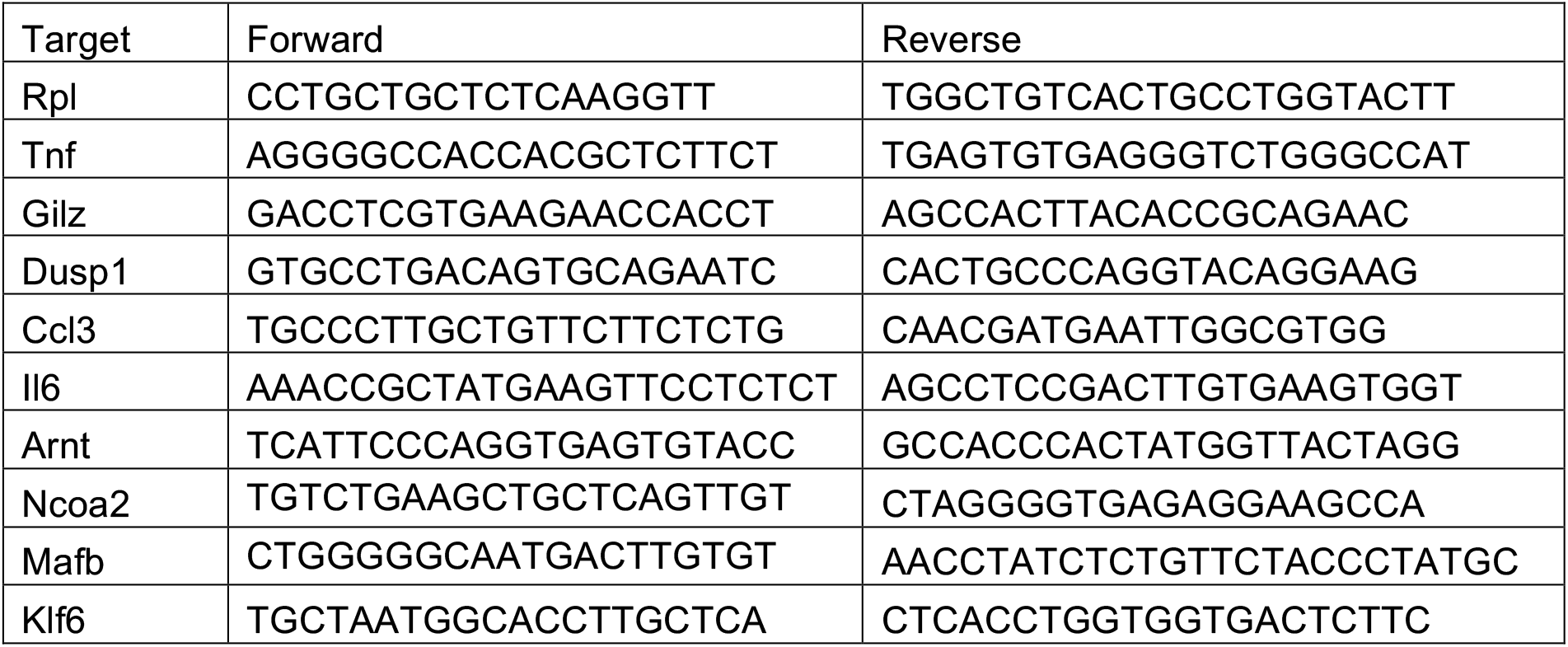

### Myoblast - macrophage coculture

Macrophages were seeded at 54,000 cells/cm^2^ in 12-well plates in DMEM 10% FBS. They were treated for 3 days with 1×10^−7^ or 1×10^−8^ M of Dex and washed. DMEM free serum was added for 24 hours to obtain conditioned medium. Myoblasts were obtained from TA muscles and were cultured in DMEM/F12 containing 20% FBS and 2% UltroserG (Pall Inc, 15950-017) as previously described (Mounier et al., 2013). For proliferation studies, myoblasts were seeded at 10,000 cells/cm^2^ on Matrigel (Corning, 356 231, 1:10) and incubated for 1 day with macrophage-conditioned medium + 2.5% FBS. Myoblasts were then washed, fixed and incubated with anti-Ki67 antibody (Abcam, ab15580, 1:100) visualized using cy3-conjugated secondary antibody (Jackson Immunoresearch, 711-165-152, 1:200). For differentiation experiments, myoblasts were seeded at 30,000 cells/cm^2^ on Matrigel (Corning, 356 231, 1:10) for 3 days with macrophage-conditioned medium containing 2% Horse Serum (Gibco). Cells were washed, fixed and stained with anti-desmin antibody (Abcam, ab32362, 1:200) and visualized by cy3-conjugated secondary antibody (Jackson immunoresearch, 711-165-152, 1:200). At least 9 ± 1 pictures *per* condition were acquired at x20 magnification on an Axio Imager.Z1 microscope (Zeiss) connected to a CoolSNAP MYO CCD camera (Photometrics) using MetaMorph Software. Quantification of proliferation (Ki67^pos^ cells) and of differentiation (number of myonuclei *per* cell) was done manually using ImageJ software.

### Glucocorticoid receptor subcellular localization

BMDMs were seeded at a density of 20,000 cells in 96 well-plates. Medium was changed to Macrophage-SFM (Thermofisher, 12065074) with 1% penicillin/streptomycin, 1% L-glutamine, 1% sodium pyruvate, 1% fungizone overnight. The following day, cells were treated with 100nM Dex (Sigma-Aldrich, D4902) for 1 hour, and fixed with 4% PFA for 15 minutes at 4°C. PFA was removed, and samples were washed with PBS 3 times. Cells were permeabilized with permeabilization buffer (PBS, 0.1% TritonX-100) for 30 minutes at 4°C, then non-specific binding sites blocked with blocking buffer (PBS, 0.1% TritonX-100, 1% fetal calf serum) for 1 hour at room temperature, then overnight at 4°C. Cells were incubated with anti-GR antibody (Cell Signaling D8H2 3660, 1:200) overnight at 4°C. After washing with PBS 5 times, cells were incubated with a fluorescently labelled secondary antibody (Thermofisher, A11037 anti-rabbit IgG Alexafluor 594) for 2 hours at room temperature. Cells were washed 5 more times and incubated with 1μg/mL DAPI (Sigma-Aldrich, D9542) for 10 minutes at room temperature. Cells were washed twice more and retained in PBS for immediate imaging. Images were acquired using the ImageXpress MicroConfocal (Molecular Devices), at least 12 images were acquired per biological sample. Nuclear GR intensity was determined using ImageJ.

### Western blot

AMPK/ACC: Macrophages were seeded at 208,000 cells/cm^2^in 6-well plates. Cells were treated with Dex (Sigma-Aldrich, D4902, 1×10^−7^ M or 1×10^−8^ M) and 991 (Spirochem, 129739-36-2, 1 μM) for 0.5 to 1 hour in DMEM containing 10% of charcoal stripped serum (Gibco, 12676029). Total protein extract was obtained in cell lysis buffer containing 50 mM Tris-HCL (pH 7.5), 1 mM EDTA, 1 mM EGTA, 0.27 M sucrose, 1% triton, 20 mM glycerol-2-phosphate disodium, 50 mM NaF, 0.5 mM PMSF, 1 mM benzamidine, 1 mM Na_3_VO_4_ and 1% cocktail phosphatase inhibitor 3 (Sigma-Aldrich, P0044). Thirty micrograms of protein were subjected to SDS-PAGE (100V) and transferred onto a nitrocellulose membrane overnight at 4°C, 30V. Blots were probed with antibodies against p-AMPKa (Cell Signaling, 2535, 1:1000), AMPKa (Cell Signaling, 2532, 1:1000), p-ACC (Cell Signaling, 3661, 1:1000), ACC (Cell Signaling, 3662, 1:1000), with β-actin (Sigma-Aldrich, A5316, 1:2000) as a loading control. p-AMPK*α* immunoblots and AMPKa immunoblots were performed on separated blot and each independent experiment was also performed on independent Western blot. Blots were quantified using ImageJ software and results represent (p-AMPKα/β-actin) ratio on (AMPKα/β-actin) ratio for each experiment and they are also normalized to the control condition.

GR/p38: Macrophages were seeded at confluency in 10cm plates, and changed to DMEM containing charcoal stripped serum (Sigma-Aldrich, F7524) overnight. The following day, cells were treated with vehicle (DMSO) or Dex (Sigma-Aldrich, D4902, 1×10^−7^ M) for 1 hour. Cells were then washed twice in ice cold PBS and total protein was extracted using Radio-Immunoprecipitation Assay (RIPA) buffer (50 mM Tris-HCl pH 7.4, 1% NP40, 0.25% sodium deoxycholate, 150 mM NaCl, 1 mM EDTA) containing protease inhibitors (Roche, 04694124001), phosphatase inhibitor cocktail 2 (Sigma-Aldrich, P5726) and phosphatase inhibitor cocktail 3 (Sigma-Aldrich, P0044). 25 μg of protein was subjected to SDS-PAGE (100V) and transferred to a nitrocellulose membrane for 1 hour at 4°C, 100V. Blots were blocked with 5% BSA (Sigma-Aldrich) and probed with antibodies against p-GR (Cell Signaling, 4161, 1:1000), GR (Cell Signaling, 3660, 1:1000), p-p38 (Cell Signaling, 4511, 1:1000), p38 (Cell Signaling, 8690, 1:1000) with β-actin (Sigma-Aldrich, A5316, 1:2000) or α-tubulin (Santa Cruz, sc-8035, 1:1000) as a loading control. Blots were quantified using ImageJ.

### *Ex vivo* phagocytosis assay

TA muscle was damaged as described above. Two and a half days after injury, mice were treated with i.p. injection of Dex (Sigma-Aldrich, D2915, 1 mg/kg). At day 3, TA muscles were harvested, digested for 1 hour with collagenase (Roche, 11088831001, 1 mg/ml) in RPMI medium at 37°C. Digested muscles were filtered through 30 μm filter with DMEM containing 50% FBS. CD45^pos^ cells were isolated using magnetic sorting (Milteny Biotec, 130-042-201 and 130-052-301) and were seeded at 190,000 cells *per* wells in 12-well plates in DMEM serum free. Primary myoblasts previously stained with PKH67 (Sigma-Aldrich, MINI67) were first submitted to 20 mM of H_2_O_2_ for 20 minutes to induce necrosis before being added to wells at 1:3 ratio (570,000 myoblasts *per* condition). After 6 hours of phagocytosis, cells were harvested, blocked with FcR Blocking Reagent (Miltenyi Biotec, 130-059-901) and thereafter stained with anti-CD45 antibody (eBioscience, 25-0451-82, 1:200), anti-Ly6C antibody (eBioscience, 12-5932-82, 1:400) and anti-CD64 antibody (BD Pharmingen, 558539, 1:100) for flow cytometry analysis.

### *In vitro* phagocytosis assay

BMDMs were isolated as described above. After differentiation, 100,000 BMDMs *per* well were seeded into 12 well plates, allowed to adhere and medium changed to L929 conditioned medium with charcoal stripped serum. Simultaneously, thymuses were isolated from 2 mice, and treated over-night with Dex (Sigma-Aldrich, D4902, 1×10^−6^M) in RPMI-1640 (Sigma-Aldrich, R8758) to induce apoptosis. The next day, apoptotic thymocytes were washed with cold PBS and FCS to remove excess Dex, then 7×10^6^ cells were incubated with CSFE (Thermofisher, C34554, 1:25000) for 20 minutes at 37°C. Labelled thymocytes were then added to BMDMs in a ratio of 5:1 and incubated at 37°C for 2.5 hours to allow phagocytosis. A negative control was maintained at 4°C. Cells were then washed twice with PBS, and scraped into FACS buffer (PBS, 10mM EDTA, 1% BSA). The cell mixture was first blocked with anti-FcɣRII/III (eBiosciences, 14-0161-82) then stained with anti-F4/80 (Invitrogen, BM8, 1:300) and analysed on a flow cytometer (LSRII, BD). Data were analysed with FlowJo, with signal higher than the 4°C control considered a positive phagocytosis event.

### Cytokine multiplex

Cytokines were determined in BAL fluid using a Cytokine & Chemokine Convenience 36-Plex Mouse ProcartaPlex Panel 1A (ThermoFisher) using a Bioplex-200 System (BioRad). Samples below the detection threshold were excluded from analysis.

### Transcription factor binding site analysis

Genes up- or down-regulated by DEX were analyzed for over-represented transcription factor binding sites using the Single Site Analysis function from oPOSSUM-3 software (Kwon et al., 2012). Sequences from 2000 bp upstream to 2000 bp downstream of the transcription start sites were analyzed against all the 29347 genes in the oPOSSUM database using a 0.4 conservation cut-off and a 85% matrix score threshold. Enriched transcription factor binding sites were ranked by Z-score and only the 15 highest scores were considered.

### Gene Set Enrichment Analysis

A FOXO3-dependent gene set was defined using GSE37051, comparing unstimulated wildtype BMDMs to unstimulated FOXO3 deficient BMDMs, analysed by GEO2R. Gene set enrichment was performed using GSEA 4.0.3 (Mootha et al., 2003; Subramanian et al., 2005).

### Chromatin Immunoprecipitation

ChIP was performed as described elsewhere (Nelson et al., 2006), with some modifications. 20×10^6^ BMDMs were changed to Macrophage serum free medium (Gibco) overnight before treatment with vehicle (DMSO) or dex (100nM) (Sigma) for 2 hours. Cells were washed twice with ice cold PBS then fixed for 15 minutes with 1% formaldehyde (ThermoScientific 28908). Formaldehyde was quenched using 1M Glycine (Sigma, 33226) for 5 minutes at room temperature. Cells were then washed twice more with ice cold PBS before homoengisation with a Dounce homogenizer (Active Motif, 40415) in Fast IP buffer (150mM NaCl, 50mM Tris-HCl (pH7.5), 5mM EDTA, 0.5%v/v NP-40, 1%v/v Triton X-100) with protease inhibitors (cOmplete Tablets EDTA-free, EASYpack, Roche, 04693132001). Resulting nuclei were centrifuged at 12,000 RPM and resuspended in shearing buffer (1% SDS, 10mM EDTA pH 8, 50mM Tris pH8) and sheared in 1.5ml Bioruptor Microtubes (Diagenode, C30010016) in a Bioruptor (Diagenode) for 8 cycles 30 seconds on, 30 seconds off. Resulting sheared chromatin was cleared by centrifugation for 10 minutes at 12,000 RPM, 4°C before diluting 1/10. 1ml of diluted chromatin was incubated with 3μg anti-FOXO3 antibody (Santa Cruz D-12, sc-48348), or 3μg of isotype control antibody (Rabbit DA1E mAb IgG XP Isotype Control, 3900, Cell Signaling Technologies) rotating overnight at 4°C. Chromatin-antibody mixes were then incubated with 20μL Protein A Dynabeads (Invitrogen, 10001D) for 3 hours, rotating at 4°C. After washing with Fast IP buffer, beads were eluted, and DNA precipitated. Chromatin was then analysed by qPCR and calculated as fold enrichment over IgG control.

### Statistical analysis

All experiments were performed at least using 3 independent primary cultures. For *in vivo* experiments, at least 4 mice were analyzed from three independent experiments for each group. Statistic tests were performed using GraphPad Prism software. Normality was tested with Shapiro-Wilk test. Depending on the experiment and the normality, parametric (Student *t* test, ANOVA with post-hoc comparisons) or non-parametric (Wilcoxon-Mann-Whitney, Kruskal-Wallis with post-hoc comparisons) tests were used. Gehan Breslow Wilcoxon test was used for survival experiments. *p* < 0.05 was considered significant.

---

**Figure Supplemental 1.**
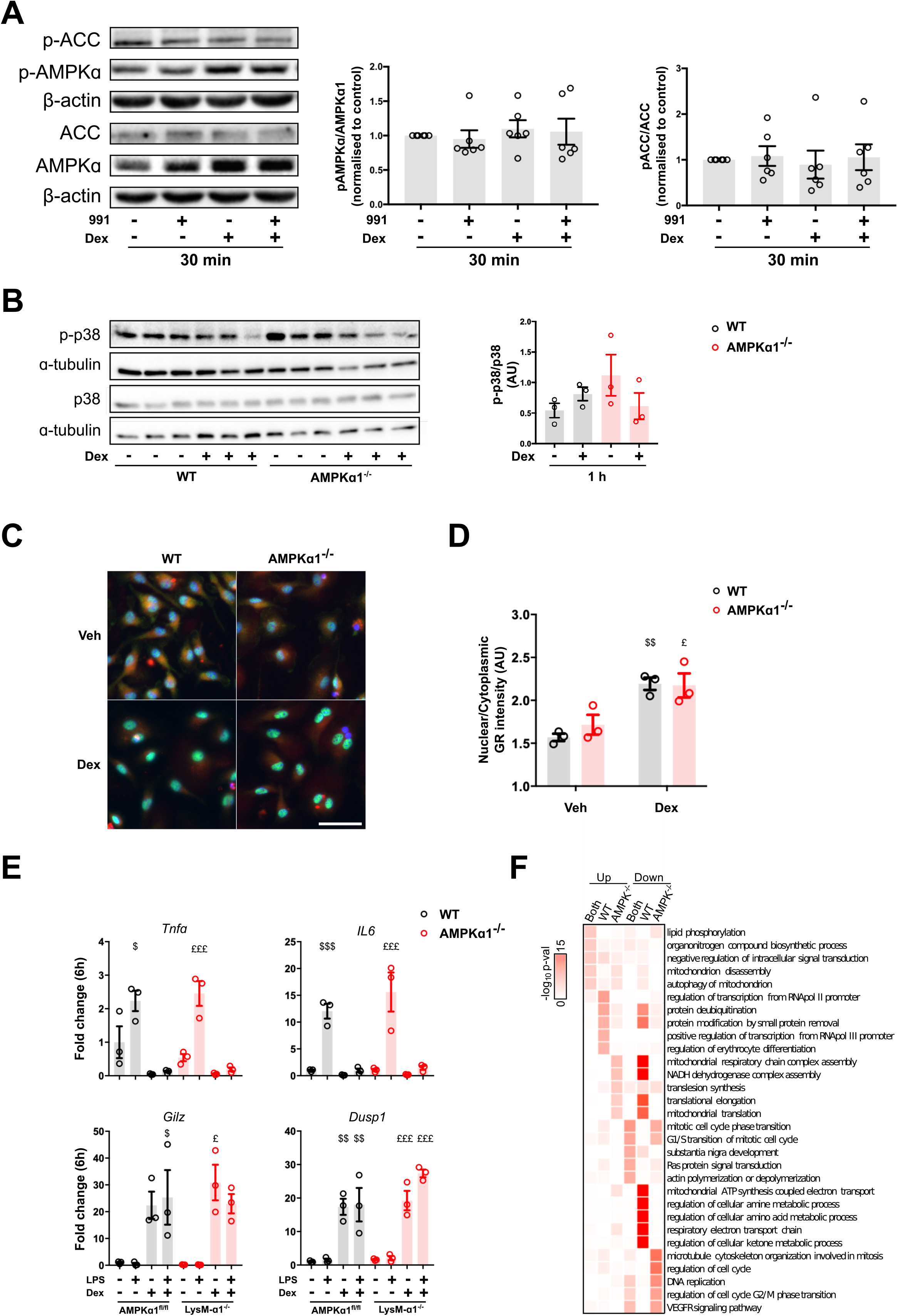
**(A)** BMDMs were treated with 991 and with Dex for 30 minutes and the phosphorylation of the AMPK targets ACC on Ser79 and that of Thr172 of AMPKα were assessed by immunoblot. Quantifications show ratio between the phospho-protein and total-protein both normalized to their respective loading control. **(B)** BMDMs from WT and AMPKα1^−/−^ mice were treated with Dex for 60 minutes and phosphorylation of p38 at Thr180/Try182 were evaluated by immunoblot. Quantification of p38 phosphorylation shows ratio between the phospho-protein on total-protein normalized to loading control. **(C-D)**WT and AMPKα1^−/−^ BMDMs were treated with Dex for 1 hour (C), and nuclear GR intensity determined by immunofluorescence was compared to cytoplasmic GR intensity (D). **(E)** RT-qPCR was performed on AMPKa1^fl/fl^ and LysMA-α1^−/−^ macrophages treated with LPS and Dex for 6 hours. (**F**) Gene ontological analysis of the AMPK-dependent transcriptome (i.e. WT selective) in response to GC treatment in WT and AMPKα1^−/−^ BMDMs. Results are means ± SEMs of 6 (A) or 3 (B-C-D) experiments. $ p<0.05, $$ p<0.01, $$$ p<0.001 *vs* untreated WT; £ p<0.05, ££ p<0.01 *vs* untreated LysM-α1^−/−^/AMPKα1^−/−^, using Student’s t test (A) or one-way ANOVA (B-D-E). Bar= 70μm (C).

**Figure Supplemental 2.**
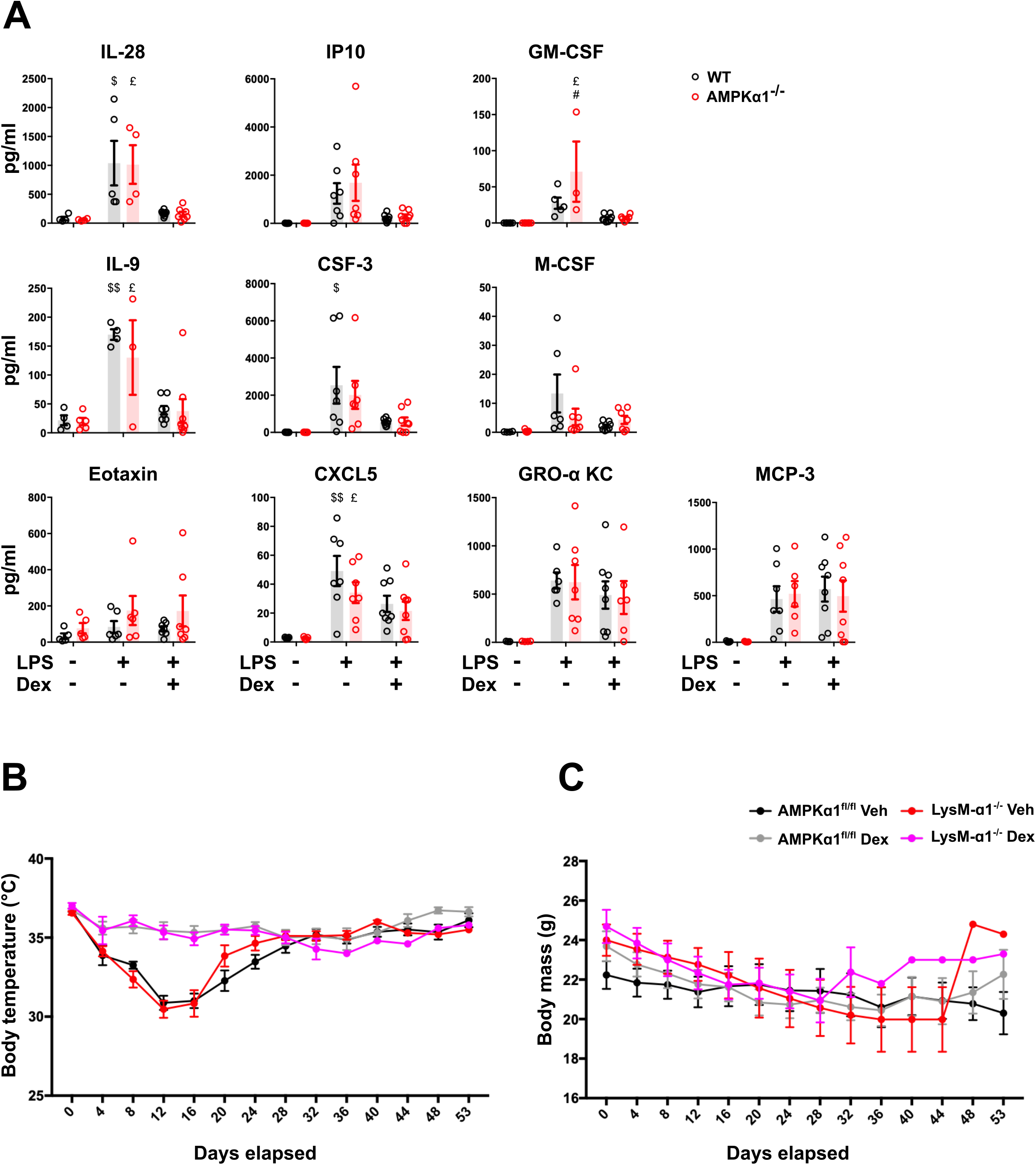
Mice were treated as described in legend of Figure 4. **(A)** The BAL cytokine content was measured by Luminex after 24 hours. (**B-C**) Graphs represent the body temperature curve (**B**) and the body mass (**C**) for each group. Results are means ± SEMs 3-8 (A), or 13-15 (B,C) animals. $ p<0.05, $$ p<0.01, *vs* untreated WT, £ p<0.05 *vs* untreated AMPKα1^−/−^; # p<0.05, compared AMPKα1^−/−^ *vs* WT for each condition, using ANOVA.

**Figure Supplemental 3.**
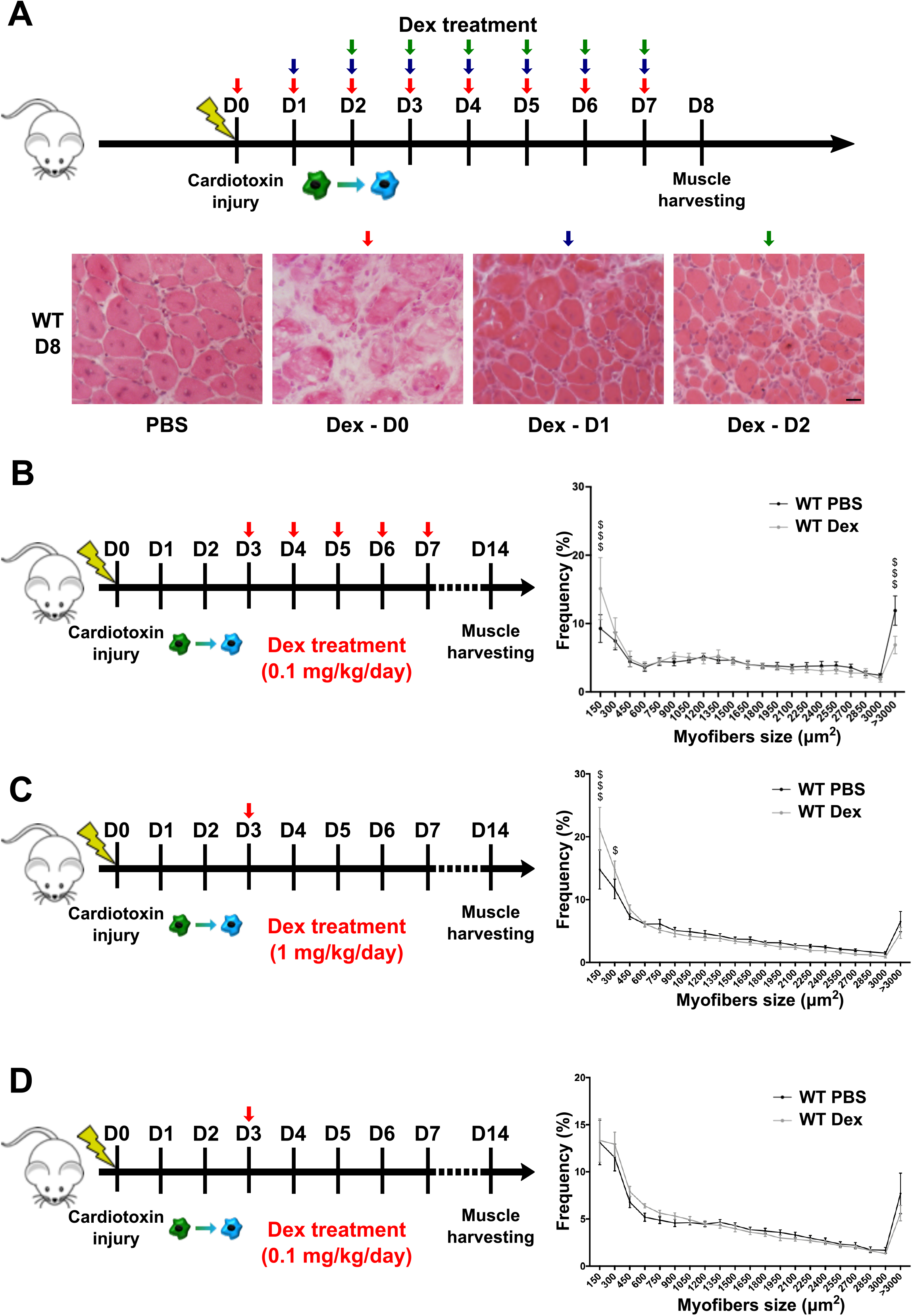
**(A)** After cardiotoxin-induced injury, WT mice were treated from day 0 (D0) to day 7 (D7) (red arrows), from day 1 (D1) to D7 (blue arrows) or from day 2 (D2) to D7 (green arrows) with Dex (i.p. 0.05 mg/kg) and *TA* muscles were harvested at day 8 (D8). Representative pictures show hematoxylin and eosin staining as compared with PBS-injected muscle (PBS). **(B-D)** After cardiotoxin-induced injury, WT mice were treated from D3 to D7 with Dex (i.p. 0.1 mg/kg) **(B)** or at D3 with Dex (i.p. 1 mg/kg) **(C)** or at D3 with Dex (0.1 mg/kg) **(D)**, and *TA* muscles were harvested at day 14 (D14) for myofiber size measurement. Graphs show the distribution of myofibers according to their size. Results are means ± SEMs of 7 (B), 8-10 (C,D) animals. $ p<0.05, $$$ p<0.001 *vs* untreated WT, using ANOVA tests.

**Figure Supplemental 4.**
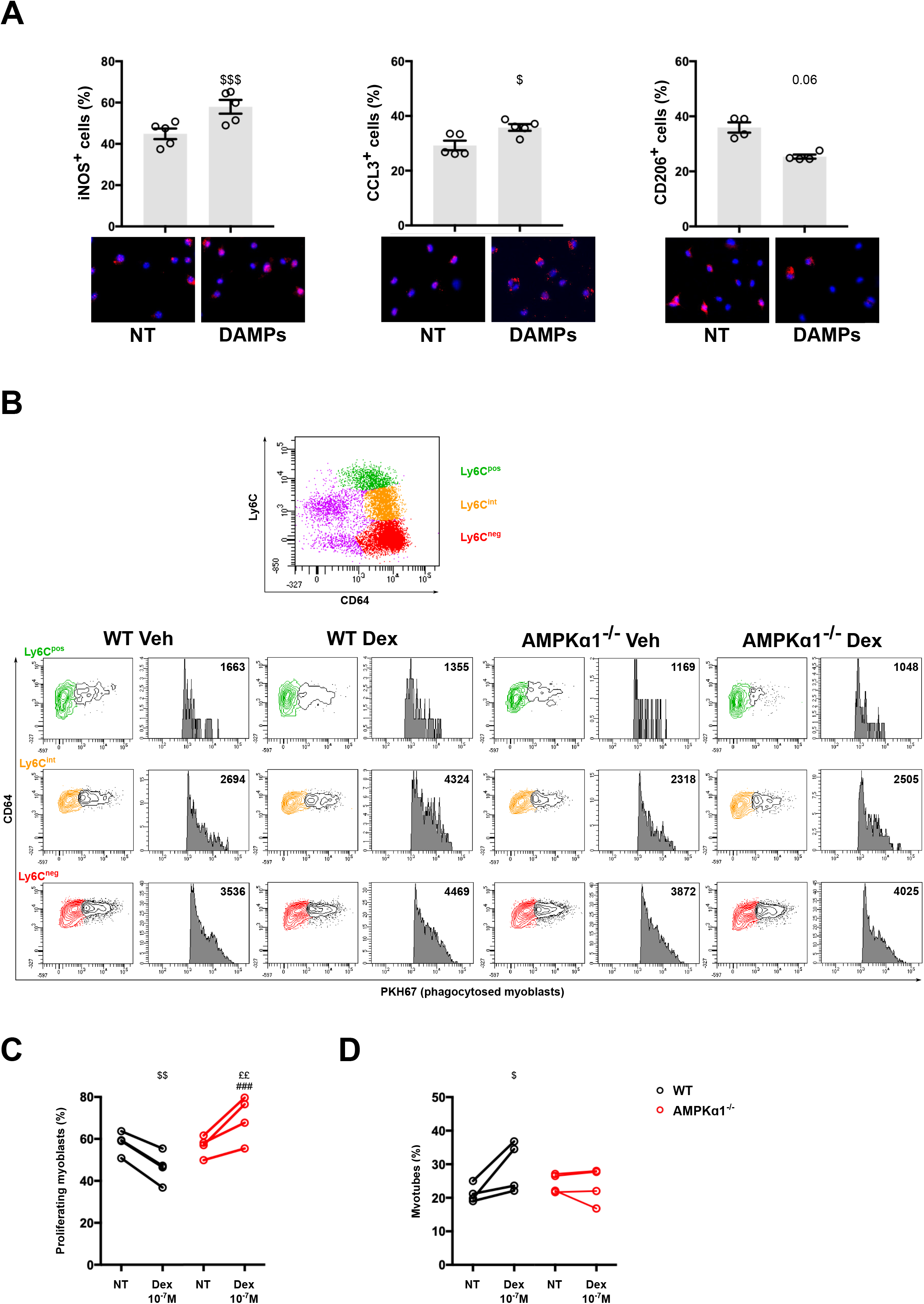
**(A)** Macrophages were treated by muscle DAMPs for 24 hours and the number of cells expressing the pro-inflammatory markers iNOS and CCL3 and the anti-inflammatory marker CD206 was measured by immunofluorescence. **(B)** Muscles and cells were treated as described in the legend of Figure 4C. Plot on top shows the whole population of macrophages stained with CD64 and Ly6C. Plots on bottom show proportion of macrophages that have phagocytosed fluorescent dead myoblasts in the Ly6C^pos^ (green population), Ly6C^int^ (orange population) and Ly6C^neg^ (red population) macrophages in the various conditions. (**C-D**) WT and AMPKα1^−/−^ BMDMs were treated with Dex (10^−7^ M) for 72 hours, washed and the conditioned medium was recovered after 24 hours and tested on muscle stem cells that were tested for their proliferation using Ki67 immunolabeling **(C)** or their differentiation using desmin labeling and counting of the number of myonuclei in the cells **(D)**. Results are means ± SEMs of 4-6 experiments. $ p<0.05, $$ p<0.001, $$$ p<0.001 *vs* untreated WT, ££ p<0.01 *vs* untreated AMPKα1^−/−^, ### p<0.001 compared AMPKα1^−/−^ *vs* WT, using Mann-Whitney-Wilcoxon test (A), ANOVA (C-D). Bar=10 μm (A).

**Figure Supplemental 5.**
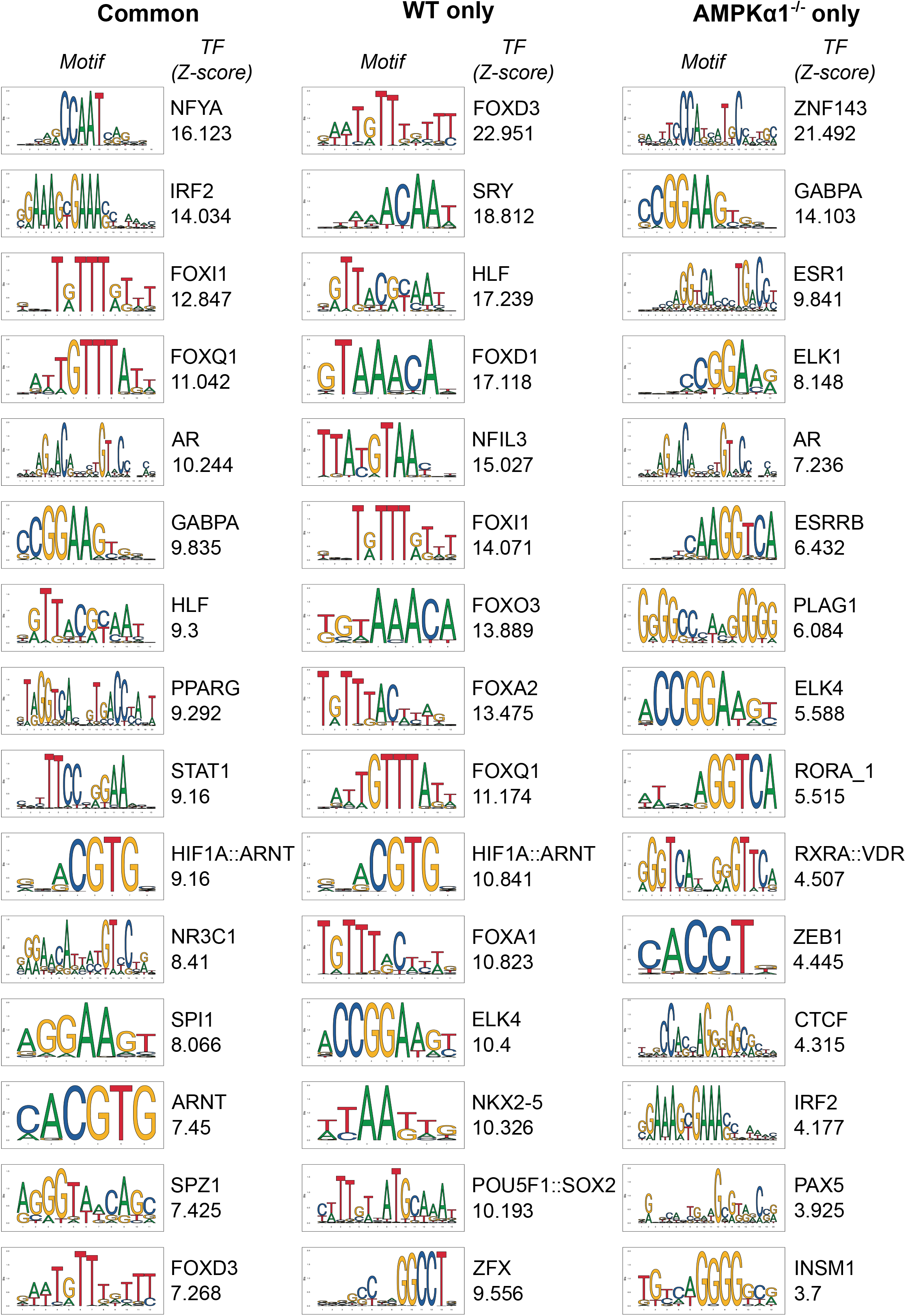
Top 15 transcription factor binding sites enriched in Dex-activated genes commonly in WT and AMPKα1^−/−^ macrophages (left panel), in WT cells only (middle panel) and in AMPKα1^−/−^ cells only (right panel), ranked by decreasing Z-score. For each transcription factor is shown its consensus binding motif from the JASPAR database and its associated Z-score.

**Figure Supplemental 6.**
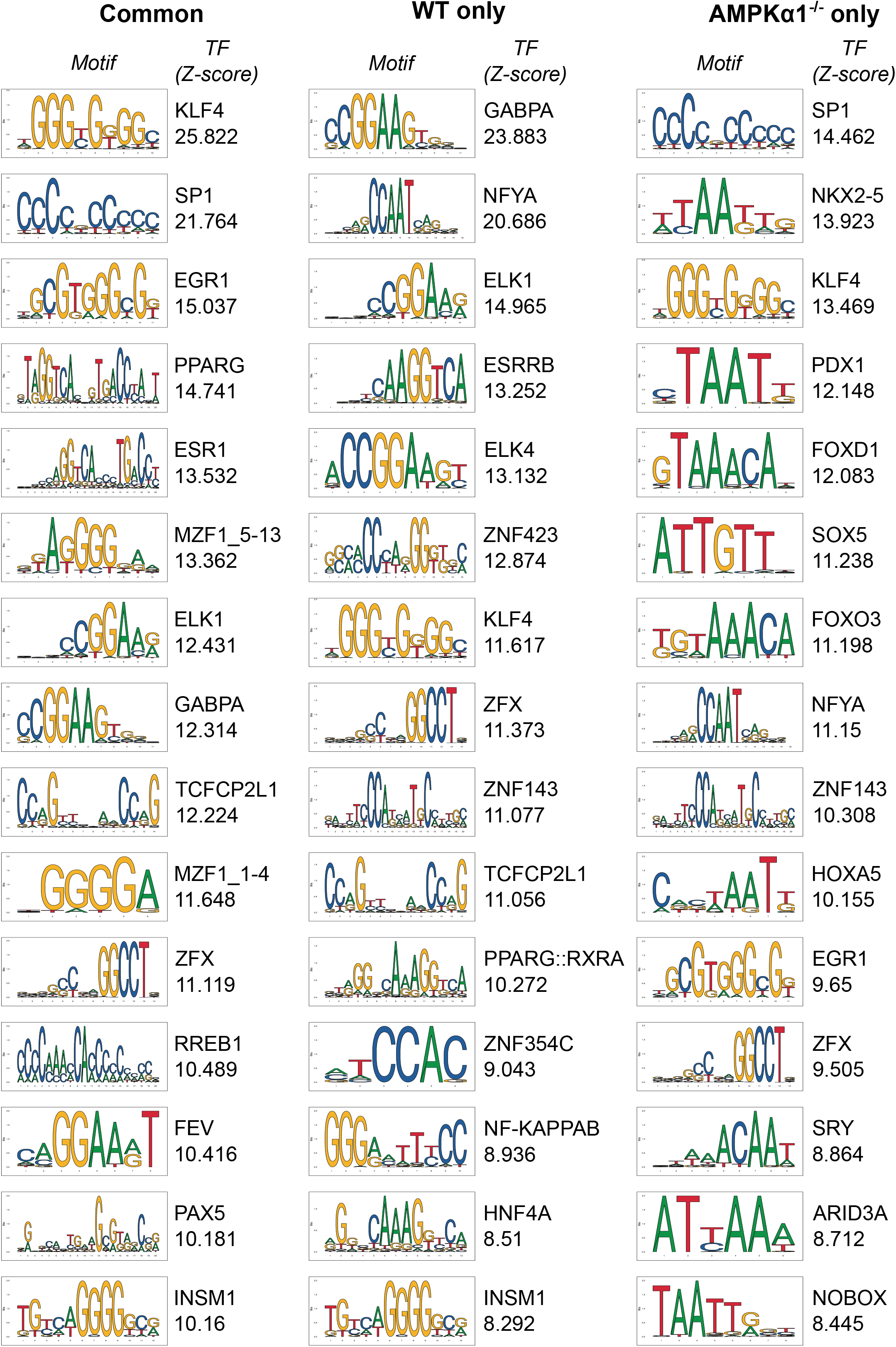
Top 15 transcription factor binding sites enriched in Dex-repressed genes commonly in WT and AMPKα1^−/−^ macrophages (left panel), in WT cells only (middle panel) and in AMPKα1^−/−^ cells only (right panel), ranked by decreasing Z-score. For each transcription factor is shown its consensus binding motif from the JASPAR database and its associated Z-score

**Figure Supplemental 7.**
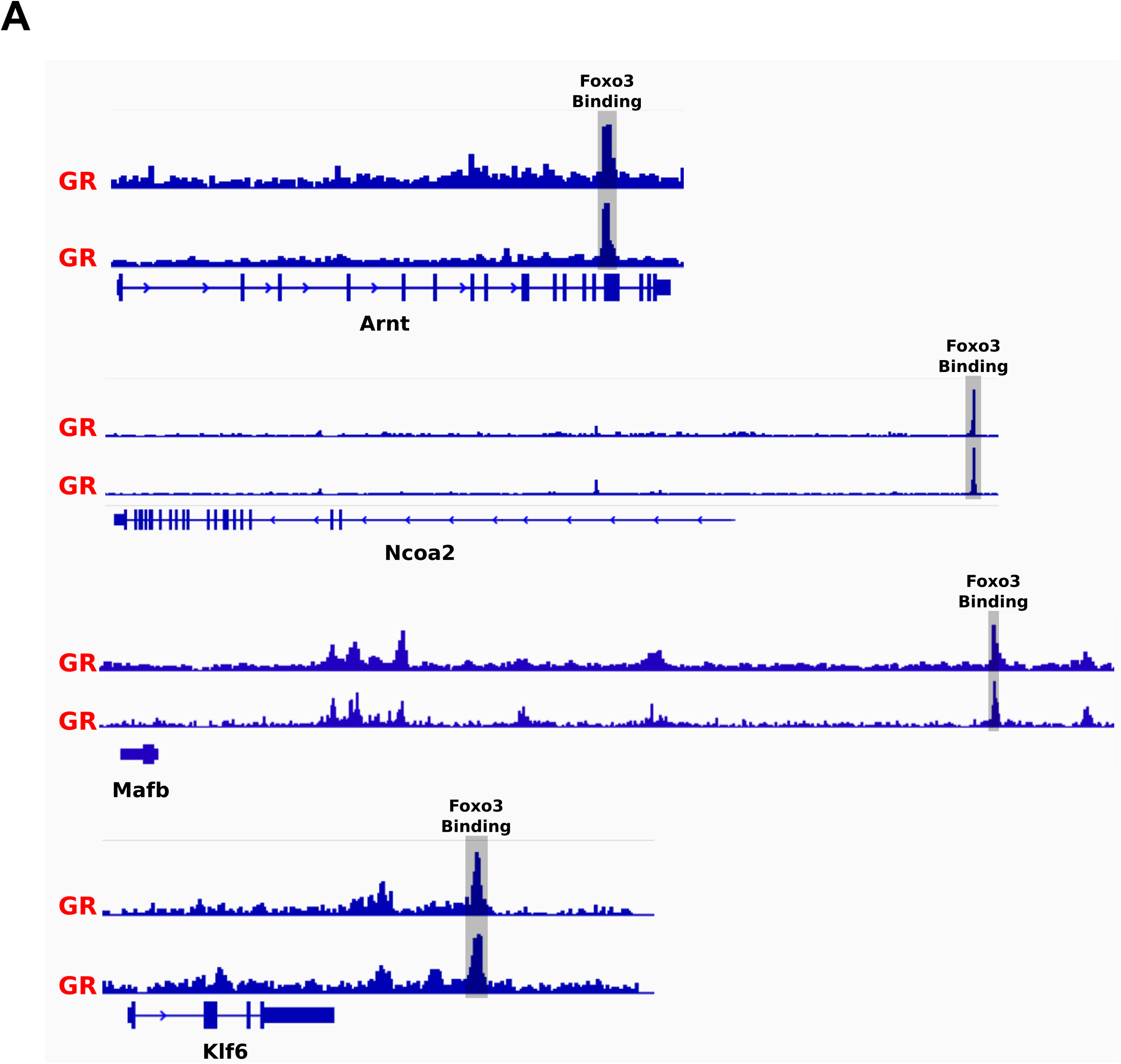
GR binding sites at WT selective genes were used as sites to look for FOXO3 binding using publically available ChIP-seq data. Gray box indicates sequence used to generate primers for ChIP-PCR.

